# Internal checkpoint regulates T cell neoantigen reactivity and susceptibility to PD1 blockade

**DOI:** 10.1101/2020.09.24.306571

**Authors:** Douglas C. Palmer, Beau R. Webber, Yogin Patel, Matthew J. Johnson, Christine M. Kariya, Walker S. Lahr, Maria R. Parkhurst, Jared J Gartner, Todd D Prickett, Frank J. Lowery, Rigel J. Kishton, Devikala Gurusamy, Zulmarie Franco, Suman K. Vodnala, Miechaleen D. Diers, Natalie K. Wolf, Nicholas J. Slipek, David H. McKenna, Darin Sumstad, Lydia Viney, Tom Henley, Tilmann Bürckstümmer, Oliver Baker, Ying Hu, Chunhua Yan, Daoud Meerzaman, Kartik Padhan, Winnie Lo, Parisa Malekzadeh, Li Jia, Drew C. Deniger, Shashank J. Patel, Paul F. Robbins, R. Scott McIvor, Modassir Choudhry, Steven A. Rosenberg, Branden S. Moriarity, Nicholas P. Restifo

**Affiliations:** Surgery Branch, National Cancer Institute (NCI), National Institutes of Health, Bethesda, Maryland, USA; Department of Pediatrics, University of Minnesota, Minneapolis, MN, USA; Masonic Cancer Center, University of Minnesota, Minneapolis, MN, USA; Center for Genome Engineering, University of Minnesota, Minneapolis, MN, USA; Department of Laboratory Medicine and Pathology, University of Minnesota, Minneapolis, Minnesota, USA; Molecular and Cellular Therapeutics, University of Minnesota, Saint Paul, Minnesota, USA; Intima Bioscience, Inc. New York, NY, USA; The Center for Biomedical Informatics and Information Technology (CBIIT), National Institutes of Health, Bethesda, Maryland, USA; National Institute of Allergy and Infectious Disease (NIAID), National Institutes of Health, Bethesda, Maryland, USA; Department of Genetics, Cell Biology and Development, University of Minnesota, MN, USA

**Keywords:** CRISPR, Checkpoint, scRNAseq, Neoantigen, TIL, PD-1, Immunotherapy, T Cell Therapy, Cancer, CISH

## Abstract

While neoantigen-specific tumor infiltrating lymphocytes (TIL) can be derived from in antigen-expressing tumors, their adoptive transfer fails to consistently elicit durable tumor regression. There has been much focus on the role of activation/exhaustion markers such as PD1, CD39 and TOX in TIL senescence. We found these markers were inversely expressed to Cytokine-Induced SH2 protein (CISH), a negative regulator of TCR signaling and tumor immunity in mice. To evaluate the physiological role of CISH in human TIL we developed a high-efficiency CRIPSR-based method to knock out CISH in fully mature TIL. CISH KO resulted in increased T cell receptor (TCR) avidity, tumor cytolysis and neoantigen recognition. CISH expression in the tumor resections correlated with TIL inactivity against p53 hotspot mutations and CISH KO in TIL unmasked reactivity against these universal neoantigens. While CISH KO resulted in T cell hyperactivation and expansion it did not alter maturation, perhaps by preferential PLCγ-1 and not AKT inhibition. Lastly, *CISH* KO in T cells increased PD1 expression and the adoptive transfer of *Cish* KO T cells synergistically combines with PD1 antibody blockade resulting in durable tumor regression and survival in a preclinical animal model. These data offer new insights into the regulation of neoantigen recognition, expression of activation/exhaustion markers, and functional/maturation signals in tumor-specific T cells.

## Main

T cells play a prominent role in immune-mediated cancer clearance by targeting tumor cells for destruction through recognition of their T cell receptors (TCRs)^1^. Antigen selection is critical, with on-target, off-tumor cytotoxicity resulting in detrimental autoimmunity^1^. Antigens created by somatic mutation—referred to as neoantigens—are appealing as they are expressed specifically by tumor and not somatic tissues. In addition, clinical responses have been associated with higher mutational load in the tumor, attributed to increased frequency of mutation reactive of T cells^2-6^. Neoantigen targeting has been facilitated by next-generation sequencing technologies that allow rapid identification of cognate non-synonymous point mutations in the genome of individual patient tumors^7^. Candidate neoantigens are subsequently loaded onto MHC of autologous antigen presenting cells (APCs) via tandem minigene (TMG) or synthetic peptide pools and used to screen TIL populations derived from resected tumor fragments^8^ or PBL^9^. Neoantigen-specific TIL identified in this manner have been attributed to objective clinical responses in a subset of patients with advanced epithelial cancers^10,11^, however achieving consistent clinical success remains a challenge.

The tumor microenvironment and defects in T cell signaling have been attributed to curtailed immune responses to cancer^12^. Substantial effort has been devoted to identifying and understand factors associated with immune suppression and to devise strategies to overcome them. Markers of T cell activation and exhaustion like PD1, CD39 and TOX^13^ have been associated with curtailed immunity to cancer. Recognizing these fundamental limitations prompted us and others^14^ to evaluate intrinsic factors that directly inhibit T cell recognition and destruction of tumor cells. We found that CISH (Cytokine-induced SH2 protein), an inhibitor of T cell receptor signaling and tumor immunity in mice^15^, was inversely expressed with these markers of activation/exhaustion. While CISH has been associated with susceptibility to multiple chronic infections in humans^16^ its role in human immunity against cancer remains is unclear.

Based on this differential expression we sought to characterize the molecular pathways and effector program governed by *CISH* in human T cells. We developed a clinical scale, cGMP-compliant manufacturing process for highly efficient and precise CRISPR/Cas9 *CISH* KO in human T cells and TIL. *CISH* KO in neoantigen selected TIL enhanced cytokine polyfunctionality, cytolysis and reactivity against identified neoantigens. CISH KO also unmasked functional reactivity to nonresponsive common *TP53* neoantigens. *CISH KO* enhanced the proliferation but not maturation of TIL and dissection of signal transduction pathways reveals that CISH inhibits PLC-γ1 but not AKT in response to TCR ligation. CISH deletion unleashes an activation and metabolic program that results in the up regulation of activation markers including PD1. Combination therapy of Cish KO and PD1 blockade resulted in pronounced synergistic cures in a preclinical mouse tumor model. These findings highlight the role of internal checkpoints in regulating neoantigen reactivity susceptibility to external checkpoint blockade.

## Results

### CISH expression and inducibility in effector T cells

We sought to explore the relationship between PD1 and other known markers of activation/exhaustion in TIL. We performed scRNAseq using T cell-enriched fresh TIL obtained from 7 treatment-naive melanoma patients. Combining all samples, TSNE clustering revealed *CD3* gene expression was consistent throughout (**Fig. 1a**). Using visual analysis, it appeared that activation markers such as *IFNG, PDCD1* (PD1), *TNFRSF9* (41BB), and *HAVCR2* (TIM3) were concomitantly enriched. Interestingly these markers appeared to not be present in CISH-positive clusters. We segregated T cells into *CISH*-high (above 50% median), *CISH*-low (below 50% median) and evaluated the expression of other activation markers within these groups. CISH-high T cells expressed significantly lower levels of *IFNG, PDCD1, TNFRSF9 and HAVCR2*, while CISH-low T cells expressed high levels of these activation/exhaustion markers (**Fig. 1b**). Thus, TIL isolated directly from patient tumors can be segregated into both CISH^high^PD1^low^41BB^low^TIM3^low^ and CISH^low^PD1^high^41BB^high^TIM3^high^ populations. In subsequent analysis using a cluster-based method^17^, consistent with previous reports^13,18,19^ we observed a significant positive correlation between PD1 (PDCD1) with activation/exhaustion markers TOX^20^ and CD39 (ENTPD1) and a negative correlation with the memory marker TCF7^21^(**Fig. 1c**). In contrast, CISH expression was negatively associated with TOX and CD39. While there was no significant correlation between CISH and TCF7 expression, we did observe a strong negative relationship between CISH and PD1 RNA expression.

**Fig. 1.**
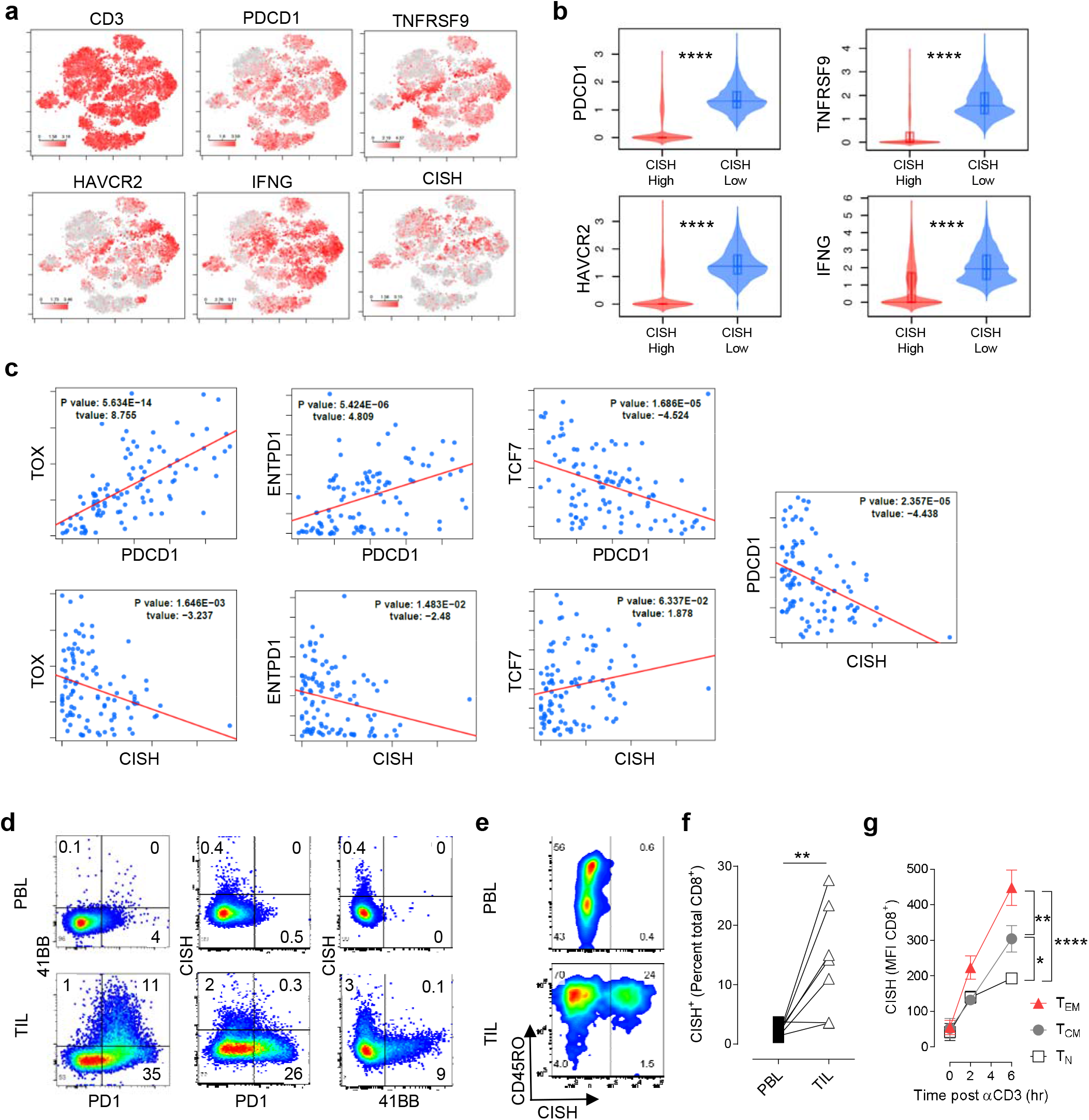
CISH is inversely expressed with markers of activation/exhaustion in melanoma patient-derived TILs. **a-c**, scRNAseq from tumor-derived T cells reveals unique clustering of CISH with other known effector/exhaustion markers. T cells were enriched from fresh tumor resections and subjected to scRNAseq (n=7 biological replicates). **b**, Violin plots of effector genes from median CISH high (>50%) of median CISH low (<50%) T cells from scRNAseq. **c**, Cluster analysis of TIL scRNAseq reveals CISH is expression is inverse to key markers of activation/exhaustion. T-value indicates positive (>1) or negative (<1) correlation. **d**, Differential expression of CISH, PD1 and 41BB in TIL. Fresh tumor resections and patient matched PBL were evaluated for co-expression of CISH, PD1 and 41BB on CD3^+^ T cells by flow cytometry. Representative of 3 patients. **e**, Flowcytometric evaluation of CISH expression in CD8^+^ T cells from PBL or matched tumor resections. **f**, Summary of increased expression of CISH in T cells from TIL tumors relative to matched PBL. n = 7 melanoma patients. **g**, Increased induction of CISH after TCR stimulation in naive CD8^+^ T cells (N), Central Memory CD8+ T (T_CM_), Effector Memory CD8^+^ T cells (T_EM_). n = 4 healthy donors. T_N_, Naïve (CD62L^+^CD45RO^-^); T_CM_, Central Memory (CD62^+^CD45RO^+^); T_EM_, Effector Memory (CD62L^-^CD45RO^+^). Statistical significance was determined by student *t* test, **P>0.01. Error bars denote +/-SEM.

To evaluate if protein expression would concur with these findings, fresh tumor fragments were obtained from three donors with matched pre-treatment PBL and evaluated for the expression of PD1, 41BB and CISH. T cells derived from PBL had low levels of 41BB, PD1 and CISH. Consistent with our scRNAseq data and previously published reports^22^, we observed that TIL derived from fresh tumors concurrently express both 41BB and PD1 (**Fig. 1d**). In concert with the scRNAseq data, the pattern of *CISH* and *PD1* expression appeared to be mutually exclusive. This differential expression pattern was also observed for CISH and 41BB. To quantify the frequency of CISH expression in TIL, minimally processed tumor resections and patient-matched pretreatment PBL were evaluated for CD8^+^ T cells and intracellular CISH expression. We observed CD8^+^ T cells from PBL had low CISH staining regardless of CD45RO status (**Fig. 1d**). In contrast to PBL-derived CD8^+^ T cells, we found a significant enrichment of CISH expressing CD8^+^ T cells in TIL directly from multiple cancer patients (**Fig. 1e**). The lack of CISH expression in CD45RO^-^ or CD45RO^+^ PBL derived T cells prompted us evaluate CISH expression and inducibility in different CD8^+^ T cells subsets. Prior to TCR stimulation, pre-sorted Naïve (T_N_), Central Memory (T_CM_) and Effector Memory (T_EM_) expressed low levels of CISH. Upon TCR stimulation a significant increase in CISH protein levels was observed, with maximal CISH levels increasing with T cell differentiation status (**Fig. 1f**). TIL are exposed to chronic antigen stimulation and typically skew toward T_CM_ and T_EM_ subsets which may account for the increased expression of CISH in CD8^+^ TIL. These data indicate in unmanipulated TIL, CISH appears to be inversely expressed with markers of an exhaustive phenotype and that CISH is not expressed in resting T cells from PBL but is readily induced by T cell activation in a manner that is linked to differentiation status. We sought to further investigate the functional significance of *CISH* in human TIL.

### Feasibility and functionality of CISH deletion in human T cells

To evaluate *CISH* function in human TIL we developed a CRISPR/Cas9-based strategy to knockout *CISH* in human T cells. We identified 3 guide RNAs (gRNA) that significantly reduced CISH protein relative to controls (**Extended Data. Fig. 1a**). Following TCR stimulation using a CD3-specific antibody, we observed increased IFN-γ production in cells edited with the 3 functional gRNAs, confirming that *CISH* KO specifically enhanced functional activation after TCR stimulation (**Extended Data. Fig. 1b**). Based on common targeting of alternate *CISH* splice isoforms and *in silico* analysis of predicted off target activity we selected a specific *CISH* gRNA targeting exon 3 for further evaluation. To characterize the immunological consequences of *CISH* KO, we manufactured *CISH* KO PBL T cells from three healthy donors, achieving an average reduction of CISH protein levels of ∼80% (**Fig. 2a**). No alteration in CISH protein expression was observed when targeting the intronic safe harbor locus *AAVS1*, confirming that *CISH* disruption was specific (**Fig. 2a**). *CISH* KO using CRISPR/Cas9 resulted in increased IFN-γ production (**Fig. 2b**) and dramatically increased TNF-α and IL-2, resulting in significantly more polyfunctional T cells after CD3-crosslinking (**Fig. 2c**). We next evaluated the role of *CISH* KO in an antigen specific response. *CISH* KO T cells were engineered with an HLA-A2 restricted NY-ESO-1-specific TCR and evaluated for cytokine production upon coculture with NY-ESO-1 positive (624) and negative (526) tumor lines. Whereas only 8% of control T cells were IFN-γ^+^ TNF-α^+^ after co-culture, 45% of *CISH* KO T cells were IFN-γ^+^ TNF-α^+^ (**Fig. 2d**). Importantly, after CISH KO there was no increase in reactivity against NY-ESO-1 negative targets, and no difference between control and *CISH* KO T cells when TCR signaling was bypassed using PMA/Ionomycin, indicating that CISH specifically inhibits TCR signaling in an antigen-specific manner. We next explored whether the increased polyfunctionality of *CISH* KO T cells enhanced tumor killing. Using a real-time tumor killing assay in relevant (TC624) and irrelevant (TC526) tumors, we observed a significant increase in Caspase3 activity in antigen-relevant tumors in the presence of *CISH* KO T cells (**Fig. 2e**). *CISH* KO did not alter response to antigen irrelevant tumors, confirming that the effect of the *CISH* KO is restricted to antigen specific recognition via TCR signaling.

**Fig. 2.**
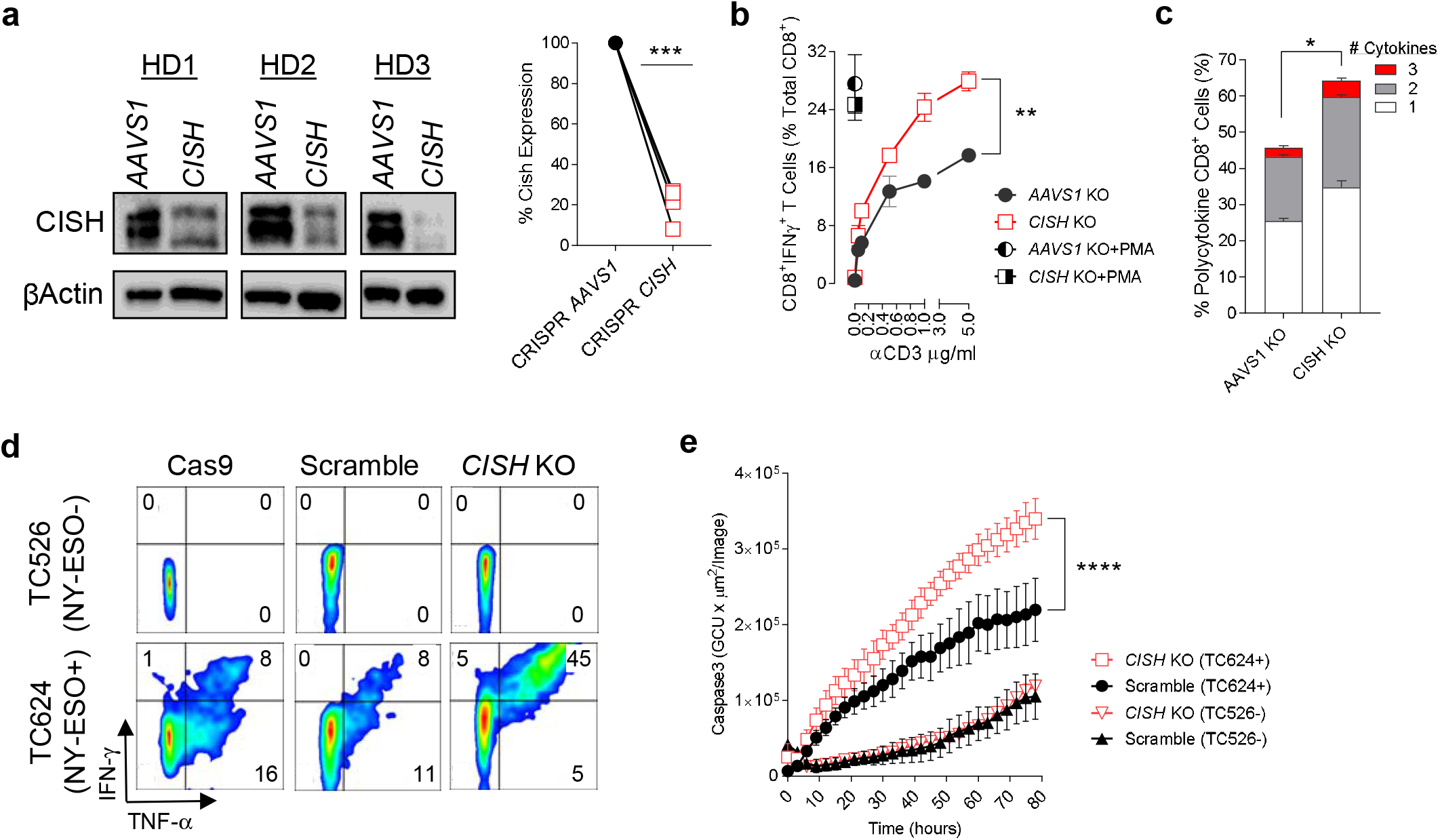
Efficient Knockout of *CISH* and hyperactivation in mature T cells. **a**, Significant reduction of CISH protein in T cells from 3 donors after CRISPR by Western blot densitometry relative to βactin. **b**, Detection of IFN-γ in CD8^+^ T cells by intracellular cytokine staining (ICS) after stimulation with titrated αCD3. **c**, Enhanced polyfunctionality after CISH deletion. Evaluation of IFN-γ, TNF-α and IL2 after CD3 stimulation and ICS. **d**, Significant enhancement in tumor reactivity after CISH deletion. Co-culture and ICS of NY-ESO-1-specific TCR transduced T cells and CRISPR of CISH with tumors expressing NY-ESO-1 (TC624) or without (TC526). **e**, Significant increase in tumor cell killing by CISH knockout T cells. Live tumor killing from (**d**) using activation of caspase3 substrate and live tumor imagining over time. (**b-d**), representative of 3 independent donors, (**e**) representative of two donors. Statistical significance was determined by either student *t* test or ANOVA for repeated measures, *P >0.05, **P>0.01, ***P>0.001, ****P>0.0001. Error bars denote +/-SEM.

### CISH knockout enhances neoantigen reactivity in TIL

The adoptive transfer of neoantigen specific T cells in patients with metastatic cancer can result in profound clinical responses^11,23^. However, identifying neoantigen reactive TIL is difficult and even when reactive TIL are identified, they often have weak reactivities that are lost after expansion. The difficulties in detection, function, and expansion of neoantigen-specific T cells may be due to the tolerogenic state of these fully mature TIL. We hypothesized that *CISH* KO may reverse this tolerogenic state and enhance TIL neoantigen reactivity. Consequently, we sought to establish a translatable process for *CISH KO* in neoantigen specific TIL (**Fig. 3a)**. To evaluate the feasibility and safety of *CISH KO* in TIL, we adapted and optimized a CRISPR-based KO strategy specifically for TIL engineering at clinical scale and in compliance with cGMP guidelines (**Extended Data Fig. 2a**). Rapid expansion protocols (REP) combining anti-CD3 stimulation and PBMC co-culture allow robust expansion of TIL^24^. Prior efforts to incorporate nuclease-based editing into the REP carried out electroporation midway through the procedure. This strategy necessitates electroporation of large numbers of TIL (>1e9) and therefore requires large amounts of editing reagent. It also involves the electroporation of a TIL/PBMC co-culture which can reduce the amount of reagent delivered specifically to TIL. As lymphocyte activation enhances nucleic acid delivery and genome editing^25^, we modified the standard REP procedure by incorporating an initial feeder-free activation step using plate-bound anti-CD3 and soluble anti-CD28 prior to electroporation (**Extended Data Fig. 2a**) Following electroporation, the TIL are transferred to PBMC feeder co-culture for REP. Using this approach, *CISH* KO in human TIL averaged >95% efficiency as measured by sequencing and TIDE^15^ analysis (**Extended Data Fig. 2b**). CISH protein loss correlated well with indel frequency at the targeted locus, averaging >95% protein reduction across the three donors (**Extended Data Fig. 2c**). Post-editing TIL expansion during the REP phase was similar to unmanipulated controls, achieving an average of >1000-fold expansion (**Extended Data Fig. 2d**).

**Fig. 3.**
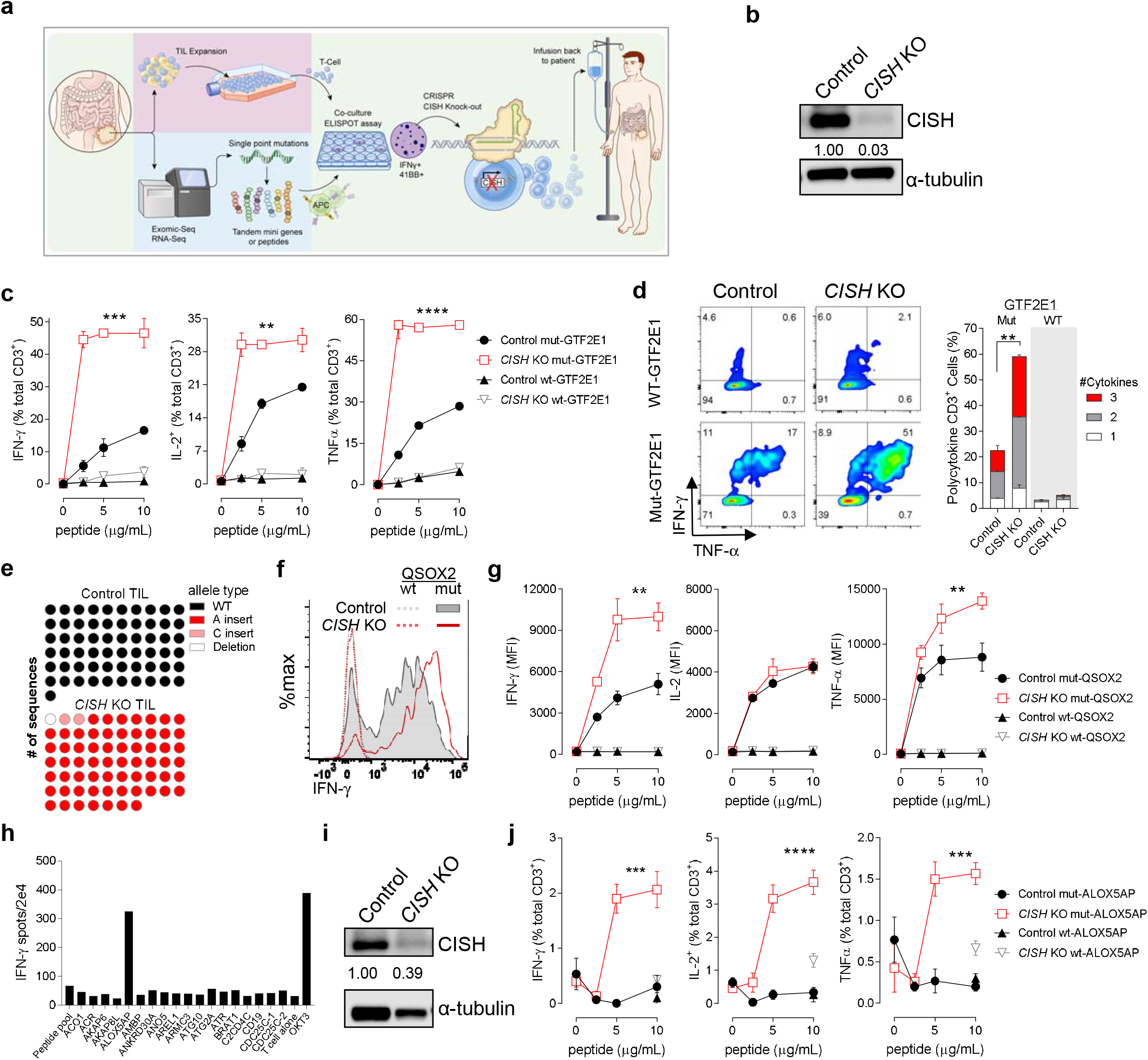
Enhanced neoantigen reactivity after the deletion of CISH in TIL. **a**, Schematic for GMP production of CISH deficient TIL. The tumor is split after excision. In one fragment T cells are grown out and in the other fragment the tumor is subjected to exomic and RNA sequencing. Non-synonymous point-mutations predicted to bind to autologous MHC are made into tandem mini genes or peptides targets. T cells derived from the tumor resection and neoantigen loaded autologous APCs are then co-cultured and assayed for the upregulation of IFN-γ or 41BB. Reactive wells are then stimulated and subjected to CRISPR mediated knockout of CISH. **b**, patient A, efficient knockout of *CISH* post CRISPR in mature TIL and evaluated by immunoblot analysis. **c**, *CISH* ko increased cytokine production and polyfunctionality after co-culture with wildtype (WT) and neoantigen loaded APCs as determined by ICS in patient A. (**d**) Evaluation of polyfunctionality of neoantigen reactive T cells after deletion of CISH in T cells co-cultured with targets in patient A. Cytokines evaluated include IFN-γ, TNF-α, and IL-2. **e**, Evaluation of CRISPR-induced disruptions in the CISH locus by Sanger sequencing in patient B. All Indels detected result in alternate coding and premature termination prior to the functional SH2 and SOCS domains. **f**, CISH deletion increased intensity of cytokine production on a per cell basis after co-culture with neoantigen loaded APCs and subjected to ICS in patient B. **g**, Significant increase in MFI of IFN-γ and TNF-α against neoantigens after CISH deletion in TIL and coculture of titrated neoantigen loaded APCs in patient B. **h**, Patient C had initial neoantigen reactivity that was lost after expansion. Initial coculture of T cells from a tumor fragment with potential neoantigen loaded APCs and assayed by IFN-γ ELISpot. **i**, Immunoblot analysis for CISH following stimulation and CRISPR mediated KO of CISH. **j**, Neoantigen reactivity was “restored” after CISH deletion in TIL and cocultured with neoantigen loaded APCs and assayed by ICS. Statistical significance was determined by either student *t* test or ANOVA for repeated measures, *P >0.05, **P>0.01, ***P>0.001, ****P>0.0001. Error bars denotes +/-SEM.

Off target (OT) editing is a key concern for production of a cellular therapy^26,27^. To evaluate OT editing, we utilized computational prediction followed by targeted next generation sequencing (NGS)^16^. At 43 candidate OT sites, we did not observe editing above the limit of detection (∼0.01%) (**Extended Table 1**). We also employed unbiased identification of double-strand breaks (DSBs) enabled by sequencing (GUIDE-seq) in TIL^11^. This technique has a lower sensitivity per OT site (detecting locus modification at frequencies of approximately 1%), but is not limited to any theoretical assumptions, e.g. based on sequence homology. Using this complementary approach, we detected no OT events in either (**Extended Data. Fig. 2e**). Thus, using two independent and complementary assays, we did not detect any OT editing in TIL modified using our *CISH-*targeting CRISPR/Cas9 reagents.

We next monitored TIL growth *in vitro* and found that *CISH* KO significantly increases and extends the proliferation of TIL compared to WT controls (**Extended Data Fig. 2f**). To rule out the possibility that *CISH* KO confers the capability of cytokine-independent growth, we removed IL-2 and observed a rapid and complete loss of viable TIL in both the *CISH* KO and WT cultures (**Extended Data Fig. 2g**). Thus, despite having significantly enhanced proliferative capacity, *CISH* KO TIL remain dependent on cytokine signaling.

Having established a clinically scalable, high-efficiency *CISH* editing process with a robust safety profile, we wanted to evaluate if this manipulation would result in enhanced reactivity against neoantigens. TIL were isolated from resected tumor fragments from three independent patients, and following whole exome and RNA-sequencing, mutation reactive TIL populations were selected as previously described^28^. TIL were isolated from a tumor resection from a 36-year-old female with metastatic colon cancer, referred to as patient A. The selected TIL population was screened against individual peptides and the cognate neoantigen was identified as GTF2E1 (S334F) (**Extended Data Fig. 3**). The neoantigen reactive TIL fragment was subjected to the approach as described (**Extended Data Fig. 2a**) and evaluated for *CISH KO* and functionality against GTF2E1 (S334F) (**Fig. 3a**). Following CRISPR editing and REP, we measured CISH protein by Western blot and observed the near-complete absence of CISH protein in the CRISPR treated TIL (∼97% reduction) (**Fig. 3b**). Next, we wanted to determine if *CISH* KO would enhance TIL recognition and functionality against the GTF2E1 (S334F) neoantigen. Autologous DCs were loaded with GTF2E1 peptide and co-cultured with control or *CISH* KO TIL for 6 hours and evaluated for production of effector cytokines. We found a highly significant increase in the percentage of IFN-γ, IL-2, and TNF-α positive CD3^+^ T cells in the *CISH* KO TIL compared to controls (**Fig. 3c**). No reactivity was observed against the wildtype (WT) GTF2E1 peptide-pulsed DCs by either *CISH* KO or control TIL. Further, maximal reactivity was maintained even at low levels of GTF2E1 neoantigen, indicating that *CISH* deletion not only increases the overall functionality of TIL but also enhances their sensitivity to antigenic stimulation. This later facet may prove critical *in vivo* where many tumors express low levels of peptide loaded MHC. In addition to increased production of individual cytokines, *CISH* KO TIL also exhibited enhanced polyfunctionality (IFN-γ, TNF-α and IL-2 positive) (**Fig. 3d**). IFN-γ^+^TNF-α^+^ frequency increased from 17% to 51% in CISH KO TIL, and there was a significant increase in IFN-γ, TNF-α, and IL-2 triple positive TIL. The ability to successfully delete CISH in fully mature human TIL and dramatically increase neoantigen sensitivity and polyfunctionality in this patient propagated us to evaluate to reproducibility across patients.

To verify our approach, a tumor from a 44-year-old male with colorectal cancer, patient B, was resected and TILs were outgrown for neoantigen screening. Reactive cultures were selected and QSOX2 (R524W) was identified as the target neoantigen (**Extended Data Fig. 4a**). Further characterization revealed that reactivity was mediated predominantly by a population of CD8^+^ TIL bearing the clonotypic Vb17^+^ TCR (**Extended Data Fig. 4b**). Since there was a predominant population of neoantigen specific T cells, we sought to determine if there was an increase in the intensity of cytokine production on a cell-by-cell basis. Editing at *CISH* was highly efficient (**Fig. 3e**), and we found a pronounced increase in the frequency and intensity of IFN-γ staining in the *CISH* KO TIL compared to the control TIL against mutant-QSOX2 loaded APCs (**Fig. 3f**). By contrast, WT QSOX2 peptide did not elicit IFN-g release in either TIL group. Using titrated neoantigen peptides, we found a significant increase in specific intensity of IFN-γ and TNF-α but not IL-2 in *CISH* KO TIL compared to control TIL (**Fig. 3g**). Despite a significant increase in functionality to mutant QSOX2 by *CISH* KO TIL, reactivity to WT QSOX2 was not detectable at any peptide concentration. From these data, it appears that *CISH* KO reproducibly increases the neoantigen-specific reactivity on a cell-by-cell basis as evidenced by a qualitative increase in cytokine production and TIL polyfunctionality, without increasing reactivity to WT antigens.

**Fig. 4.**
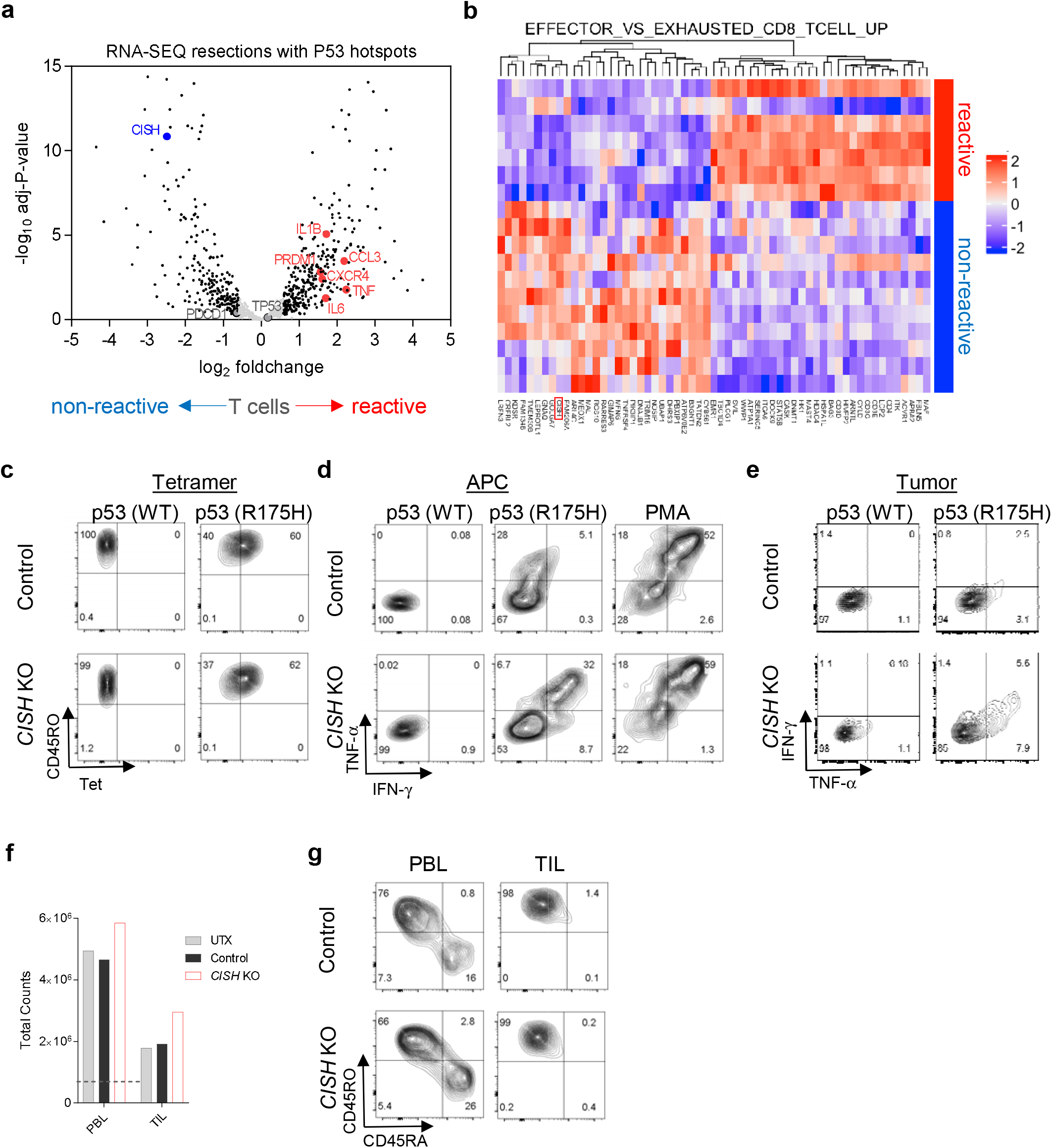
CISH KO restores TIL reactivity against universal hotspot p53 mutations. **a**, Immunological gene signature expression analysis from tumor fragments with p53 hotspot mutations that were found to have or not have detectable T cell reactivity to mutant p53. Fresh tumor fragments were sequenced for p53 hotspot mutations. 18 fragments were identified to contain known targetable p53 neoantigens. Isolated TIL from each fragment were screened for p53 neoantigen reactivity, 7 fragments were found to have p53 hotspot reactivity, while 11 did not. Volcano plot of immune-filtered genes from RNA-SEQ of tumor resections from reactive and non-reactive cultures.Gray indicates genes below 1.5-fold change and/or a p>0.05. **b**, Cluster analysis of differentially expressed immunological genes associated with effector and exhausted phenotype in CD8^+^ T cells. Each row represents an individual tumor fragment and are clustered into TIL with and without reactivity to mutant p53. **c**, WT p53 or mutant p53 (R175H) tetramer staining of TIL from patient D with or without CISH deletion. **d**, Significant increase in IFN-γ and TNF-α against p53 hotspot mutation after CISH deletion in TIL cocultured with antigen-loaded APCs and assayed by ICS. **e**, Augmented tumor reactivity after CISH deletion. Increase in IFN-γ and TNF-α staining in CISH deleted TIL cocultured with p53 hotspot mutation expressing tumors and assayed by ICS. **f**, Cell count of PBL and matched TIL from patient D after CISH deletion and 10 days of culture. Dotted line indicates initial starting count. **g**, Flow cytometric evaluation of phenotype of PBL or TIL from patient D after CISH deletion after 10 days of culture. Statistical significance was determined by either student *t* test or ANOVA for repeated measures, *P >0.05, **P>0.01, ***P>0.001, ****P>0.0001. Error bars denotes +/-SEM.

In some instances, the detection of neoantigen specific TIL in the initial screening is either lost or significantly diminished after a REP. Thus, we sought to evaluate if *CISH* KO would resurrect functionality in an instance where neoantigen reactivity was lost after a REP. To this end, we evaluated TIL from a 45-year-old female with sigmoid colon cancer, patient C, whose TIL ultimately lost neoantigen reactivity after a REP (data not shown). Following whole exome sequencing, candidate peptides were generated and pooled to test reactivity to TIL derived from fragment F9.Subsequent parsing revealed TIL to be specific to the neoantigen ALOX5AP (K153R) (**Fig. 3h**). This neoantigen-reactive TIL population was then subjected to REP with and without *CISH KO* (**Fig. 3i**). Upon co-culture with autologous APCs loaded with titrated concentrations of mutant ALOX5AP peptide, we found that control TIL had no detectable reactivity to mutated ALOX5AP peptide as measured by staining for IFN-γ, IL-2 or TNF-α (**Fig. 3j**). In contrast, we detected neoantigen reactive T cells after *CISH* KO with significant and titratable increases in IFN-γ, IL-2 and TNF-α. Moreover, WT antigen did not elicit a functional response in either group. These data indicate that loss of neoantigen reactivity after traditional TIL REP can be restored by *CISH* KO, resulting in a significant increase in TIL functionality against a tumor neoantigen.

### CISH as a determinant of neoantigen reactivity

*TP53* mutations are frequent in many cancers and can be presented as neoantigens, however only a subset of patients harbor T cells that elicit a functional immune response^29^. As CISH KO enhances and restores neoantigen reactivities in TIL, we sought to evaluate whether CISH may dampen the T cell response to common *TP53* mutations. To this end, we evaluated RNA-seq from fresh resections that contained *TP53* driver mutations and that we had tested for mutant p53 neoantigen reactivity. We evaluated 18 resections identified with TP53 driver mutations: In 7 of these, we found neoantigen reactivity (reactive), whereas we were unable to identify reactivity in 11 of these (non-reactive). To filter the heterogeneity often found within these tumor resections we applied a commonly used immunologic filter^30^. We observed delineated clustering among the TP53 neoantigen reactive and non-reactive resection fragments (**Extended Data Fig. 5a**). Volcano plot analysis displays relative gene expression, with cut off being >1.5 Log_2_ fold change and p value > 0.05 shaded in gray (**Fig. 4a**). Interestingly, *TP53* and PD1 (*PDCD1*) expression were not significantly different in either group. However, we found genes associated with T cell activation such as *TNF, IL6, PRDM1, IL1B* and chemokines *CCL3* and *CXCR4* upregulated in reactive resections. This was reflected by the enrichment of activation, differentiation, chemoattraction and signaling GSEA profiles in reactive resected fragments (**Extended Data Fig. 5b**). Conversely, *CISH* expression clustered with non-reactive fragments among genes associated with T cell activation (**Fig. 4b**). From these data it appears that *CISH* is preferentially expressed in non-reactive tumor resections, while reactive tumor resections exhibit a profile of T cell activation. Uncovering T cell reactivity to shared “hot spot” TP53 mutations would obviate the laborious process of delineating individual neoantigen TCRs for every patient^31^. The KO of CISH may improve their reactivities and make shared antigen targeting more universal. Thus, we sought to evaluate if CISH deletion would uncover reactivities from a patient with a TP53 hotspot mutation and known specificity with tetramer positive T cells.

**Fig. 5.**
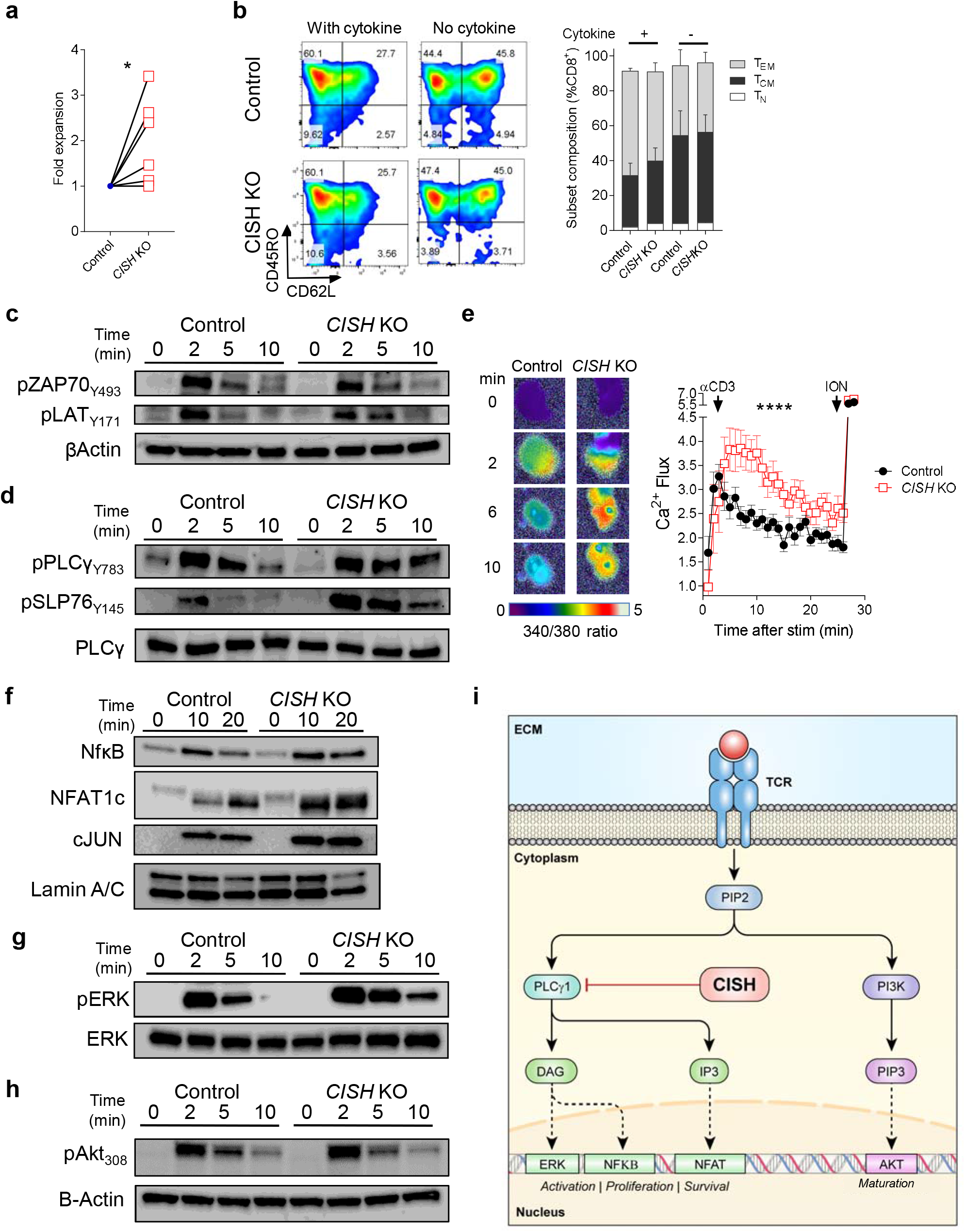
CISH KO increases T cell activation but not maturation. **a**, CISH deletion results in increased T cell expansion. Following naïve CD8^+^ T cell enrichment, T cells were TCR stimulated, CISH deleted by CRISPR mediated KO and evaluated for expansion. Relative fold-change in T cell expansion compared to non-deleted controls after 14 days of culture (*n*=7 biological replicates). **b**, CISH deletion does not alter T cell maturation. Phenotypic analysis by flow cytometry of naïve-derived CD8^+^ CISH deleted T cells with TCR stimulation and supportive cytokines or with TCR alone after 14 days of culture. No significant difference was observed in the composition of T cell subsets after CISH deletion in either condition. T_N_, Naïve (CD62L^+^CD45RO^-^); T_CM_, Central Memory (CD62^+^CD45RO^+^); T_EM_, Effector Memory (CD62L^-^CD45RO^+^). **c-h**, CISH deletions results in hyperactivation of intermediate and late TCR signaling components. Naïve-derived CD8^+^ T cells deleted for CISH and cultured for 10 days. T cells were rested overnight and were re-stimulated with cross-linked αCD3 and immunoblotted or evaluated for Ca^2+^ flux at times indicated. **c**, CISH deletion does not alter early TCR signaling. Immunoblot analysis for pZAP70_Y493_, pLAT_Y171_, and βActin at times indicated with or without CISH KO. **d**, CISH inhibits intermediate TCR signaling. Immunoblot analysis for pPLCγ1_Y783_, pSLP76_Y145_, and whole PLCγ1 after TCR ligation at times indicated with or without CISH KO. **e**, Increased Ca^2+^ flux after CISH deletion and TCR-ligation using TIRF microscopy on a lipid bilayer. Individual T cells with or without CISH deletion were followed for Ca^2+^ flux at times indicates after forming immunological synapses. (n=13-18 T cells per group). **f**, CISH deletion enhances nuclear translocation of NFκB and NFAT after TCR ligation. Immunoblot analysis for nuclear NFκB, NFAT1c, c-JUN and Lamin A/C after TCR ligation at times indicated with or without CISH KO. **g-h**, CISH deletions results in hyperactivation of ERK but not AKT. Immunoblot analysis for pERK and whole ERK after TCR ligation at times indicated with or without CISH KO. **h**, Immunoblot analysis for pAKT_S473_ and β-actin after TCR ligation at times indicated with or without CISH KO. **i**, Proposed model of how CISH may regulate T cell activation, proliferation and survival but not maturation through PLCγ1 and not AKT signaling. Representative of 3 independent donors. Statistical significance was determined by either student *t* test or ANOVA for repeated measures, *P >0.05, **P>0.01, ***P>0.001, ****P>0.0001. Error bars denotes +/-SEM.

We obtained a TIL fragment from a 36 year old female patient with metastatic colorectal cancer, patient D, that had modest neoantigen reactivity to p53 (R175H) in the context of HLA-A0201 and for which we had an estimate of precursor frequency based upon tetramer staining^32^. With a known precursor frequency, we could determine if the increase in reactivity after *CISH* KO was due to an enhancement in T cell function or simply an increased abundance of specific T cells. After *CISH* KO by CRISPR, we performed flow cytometry for p53 HLA-A0201 tetramer and evaluated TIL for reactivity to peptide and p53 (R175H) expressing tumor. We found no discernible change in tetramer staining between control and *CISH* KO T cells (∼60% of T cells in both groups) (**Fig. 4c**). Intracellular cytokine staining was performed after coculture with antigen loaded APCs (**Fig. 4d**). We found that both the control and *CISH* KO neoantigen specific T cells elicited no functional response against non-mutated p53. In the presence of mutated p53 there was a dramatic increase in IFN-γ staining from 5.4% in the control to 40.7% in *CISH* KO, with approximately a log increase in staining intensity. Similarly, there was a dramatic increase, from 5.1% to 32%, in polyfunctional cytokine production of IFN-γ^+^TNF-α^+^ after *CISH* KO. Bypassing TCR signaling using PMA/ION revealed no functional defects in the control or *CISH* KO T cells. To evaluate T cell immunity to a naturally processed and presented neoantigen, modified T cells were cocultured with HLA-matched tumors ectopically expressing full length WT or mutant p53 for 6 hours and stained for expression of intracellular cytokines (**Fig. 4e**). Like neoantigen peptide loaded APC’s, we saw an increase in cytokine production after *CISH* KO. It is important to note that while there were similar quantities of mutant p53-tetramer positive T cells in both CISH expressing and KO groups (**Fig. 4c**), only 5% of the control T cells were IFN-γ positive, and at a low MFI.

Increases in effector cytokine production typically lead to a decrease in expansion and accelerated T cell maturation^22,23^. To determine the influence of *CISH* KO on this process, we disrupted *CISH* in PBL and TIL and measured expansion and phenotype. *CISH* KO PBL and TIL exhibited a slight increase in expansion compared no electroporation (UTX) or unmanipulated controls (**Fig. 4f**). Phenotypic analysis found that PBL retained a less differentiated CD45RO^-^ CD45RA^+^ phenotype compared to TIL, and there was a slight increase in the CD45RO^-^CD45RA^+^ population in CISH KO PBL T cells (**Fig. 4g)**. These results suggest that *CISH KO* may not influence T cell maturation, although further studies are needed to better understand the apparent contradiction in hyper-effector function in the absence of hyper-maturation.

### CISH KO enhances T cell expansion and function but not maturation

PBL is a complex mixture of T cells and may not clearly represent the subtle changes in T cell maturation. Thus, we evaluated a pure population of undifferentiated T cells in which we could follow their maturation. To this end, we enriched naïve CD8^+^ T cells and studied their expansion, metabolism and maturation after stimulation with or without *CISH* KO. Here, we observed a significant increase in expansion 10 days after TCR stimulation and cytokine support of CD8^+^ *CISH* KO T cells compared to control T cells (**Fig. 5a**). CISH deletion has been shown to enhance NK cell metabolism ^33^. To evaluate if this was also true in T cells, after expansion T cells from these six donors were subjected to metabolic analysis using Ultra-HPLC-MS/MS with or without TCR stimulation. CISH KO resulted in an increase in many metabolites in glycolysis, TCA cycle and lipid metabolism which was even more pronounced 4 hours after TCR stimulation (**Extended Data Fig. 6a**). In fully mature TIL, OCR analysis revealed that CISH KO resulted in an increase in both basal and spare respiratory capacity (**Extended Data Fig. 6b**). After expansion of naive-derived CISH deleted CD8^+^ T cells we evaluated the maturation state using CD62L and CD45RO. Surprisingly, we did not observe a change in phenotypic composition in either group (**Fig. 5b**). However, cytokine support in addition to TCR stimulation can enhance T cell maturation and may mask subtle changes. While overall less mature in the absence of cytokine support, there were no overt changes in T cell maturation with or without *CISH* KO (**Fig. 5b**). This observation that *CISH* KO increases T cell functionality, killing, and expansion without influencing maturation is in stark contrast to our current understanding of the progressive maturation model^34,35^. To further probe this observation, we investigated the molecular underpinnings that could account for this divergence.

**Fig. 6.**
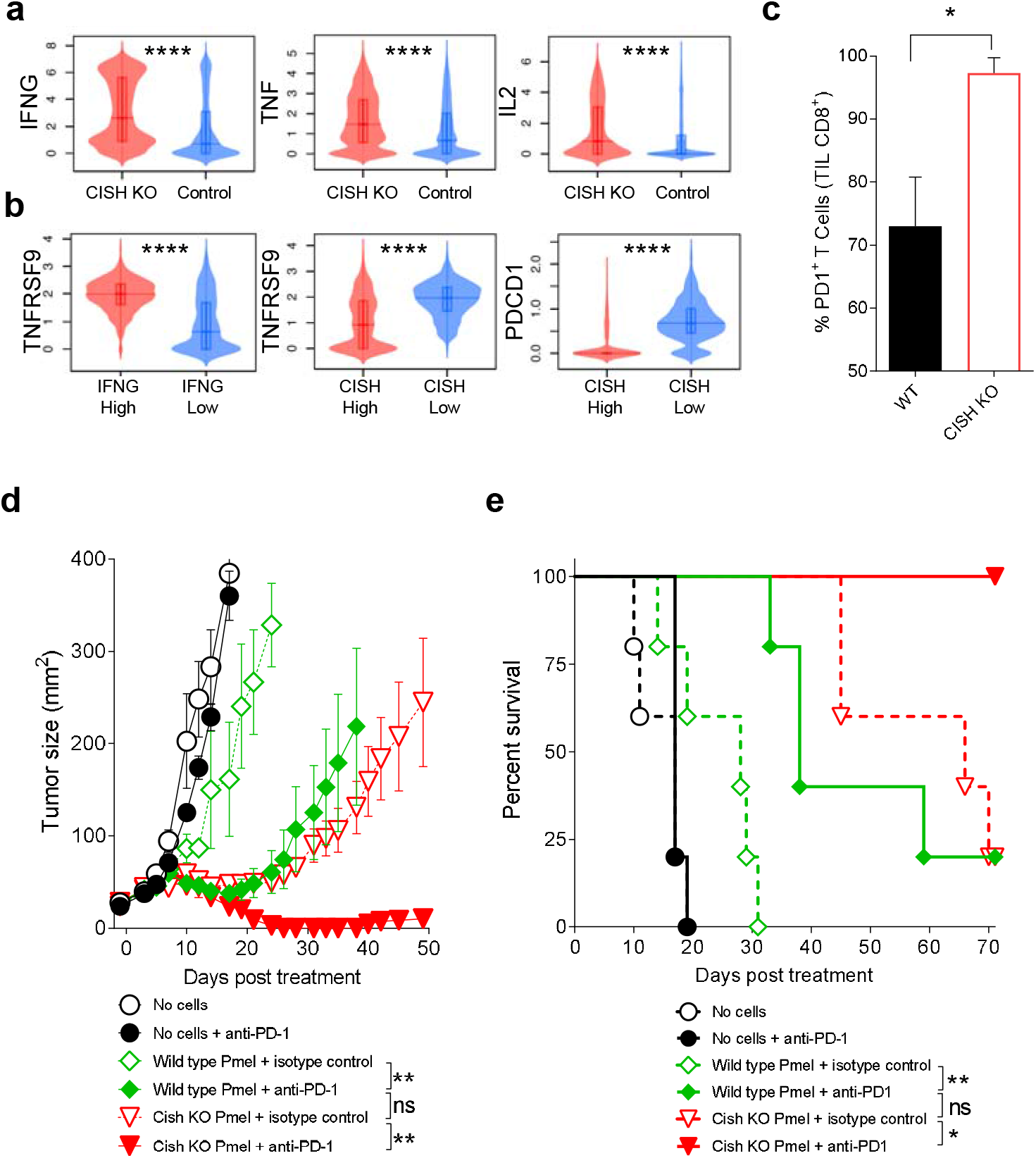
CISH KO unleashes and activation program and susceptibility to PD1 blockade. **a-b**, CISH deletion increases cytokine production and 41BB and PD1 expression. **a**, Violin plots for IFNG, TNF, IL2 gene expression from scRNAseq of T cells with or without CISH deletion 4 hours after TCR stimulation (n=3 biological replicates). **b**, Violin plots of effector genes from median IFNG or CISH high (>50%) of median IFNG or CISH low (<50%) T cells from scRNAseq of CISH deleted TCR stimulated T cells. **c**, Increased PD1 expression in TIL knocked out for CISH. Flow cytometry analysis of murine CD8^+^ TIL seven days after adoptive cell transfer (ACT) of naïve melanoma-specific T cells knocked out for CISH. **d-e**, Combination of CISH deletion and PD1 blockade significantly enhances adoptive immunotherapy. Adoptive cell transfer (ACT) of melanoma-specific T cells knocked out for Cish into B16-melanoma bearing mice with or without administration of antibodies blocking PD1 blockade. Product of perpendicular diameters blindly evaluated over time, 5 mice per group, independently repeated 3 times. **e**, Survival of mice treated in **d**. Statistical significance was determined by either student *t* test, ANOVA for repeated measures, or Log-Rank Mantel-Cox test, *P >0.05, **P>0.01, ***P>0.001, ****P>0.0001.

In mice, we observed that Cish interacts with and facilitates the degradation of the TCR signaling intermediate Phospholipase C-γ1^15,36^. Thus, we systematically evaluated the TCR signaling cascade in *CISH* KO human T cells for deviations that might account for this hyper-activation without hyper-maturation paradox. *CISH* KO CD8^+^ naïve-derived human T cells were expanded for 10 days, rested overnight in the absence of cytokine, then stimulated with TCR crosslinking and evaluated by immunoblot for early, intermediate, and late signaling factors^37,38^. TCR signaling factors ZAP70 and LAT are phosphorylated early after TCR ligation^39^, however we did not observe any changes in the phosphorylation kinetics or intensity of ZAP70 or LAT in the presence or absence of *CISH* following TCR stimulation (**Fig. 5c**). Intermediate signaling factors PLC-γ1 and SLP76 are critical in propagating the TCR signaling cascade that bridges antigen recognition with transcriptional mediators^39^. We detected a strong increase in the intensity and duration of PLC-γ1 and SLP76 phosphorylation after TCR stimulation in *CISH* KO T cells compared to control T cells (**Fig. 5d**).

Activation of the PLC/SLP76 complex results in the cleavage of phosphatidylinositol 4,5-bisphosphate (PIP2) into inositol 1,4,5-trisphosphate (IP3) and diacylglycerol (DAG). DAG activates several processes including NFκB while IP3 facilities the entry of Ca^2+^ into the cytoplasm culminating in NFAT translocation and transcriptional activation of target genes^39^. To evaluate changes in Ca^2+^ flux in T cells during TCR stimulation we used total internal reflection fluorescence microscopy (TIRFM) ^40^. We utilized an artificial lipid bilayer to evaluate TCR synapse formation during TCR stimulation and measure Ca^2+^ flux using Ca^2+^ sensitive fluorescent dyes TIRFM system. After synapse formation, individual control and *CISH* KO T cells were followed over the course of 30 minutes and evaluated for Ca^2+^ flux. We found a significant increase in the amplitude and duration of Ca^2+^ flux in CISH KO T cells after synapse formation (**Fig. 5e**). There was no difference in Ca^2+^ flux between control and *CISH* KO T cells when TCR signaling was bypassed by addition of Ionomycin (**Fig. 5e**). These data confirm that the effects of *CISH* KO are specific to TCR stimulus and are in line with the augmented levels of phosphorylated PLCγ-1 and SLP76.

Phosphorylation of PLCγ-1 and SLP76 propagates a signaling cascade culminating in the translocation of NFAT and NFκB into the nucleus and activation of the MAPK pathway^41,42^. We observed an increase in the amplitude and duration of both NFAT and NFκB translocation to the nucleus after TCR stimulation in *CISH* KO versus control T cells (**Fig. 5f**). A slight increase in c-Jun levels was also observed in *CISH* KO T cells, which is notable as increased c-Jun expression confers resistance to CAR T cell exhaustion^43^ (**Fig. 5f**). Consistent with increased TCR signaling, we observed enhanced phosphorylation of ERK in *CISH* KO T cells versus controls (**Fig. 5g**). Surprisingly, while AKT phosphorylation generally increased after TCR stimulation, there was no difference between control and hyper-activated *CISH* KO T cells (**Fig. 5h**). From these data, it appears that *CISH* KO results in increased PLCγ-1 but not AKT phosphorylation. We and others have observed that AKT activation after stimulation is largely associated with T cell matureation^44,45^. These insights offer a potential mechanism for the paradoxical enhancement of expansion and hyper-activation without associated maturation observed in *CISH* KO T cells. Thus, we propose a model where *CISH* preferentially blocks PLCγ1 signaling but AKT phosphorylation is unaffected, resulting in increased proliferation, function, and survival without altering maturation (**Fig. 5i**).

### CISH regulates expression of activation makers and susceptibility to PD1 blockade

In unmanipulated TIL we observed that CISH was inversely expressed with several activation/exhaustion markers. To evaluate the role of *CISH* in regulating the expression of these activation/exhaustion markers, we analyzed gene expression in *CISH* KO T cells by scRNAseq. *CISH* KO Naïve CD8^+^ T cells from 3 donors were cultured for a total of 10 days, rested overnight, and then TCR stimulated for 4 hours and analyzed by scRNAseq for population-based expression patterns. In concert with our previous findings, we found *CISH* KO in T cells resulted in a significant increase in *IFNG, TNF* and *IL2* gene expression (**Fig. 6a**). Not surprisingly, when T cells were grouped into *IFNG* high (above median) and low (below median), we observed high levels of *41BB* (*TNFRSF9*) in *IFNG*^high^ and not *IFNG*^low^ group (**Fig. 6b**). Conversely, when T cells were separated based upon *CISH* expression into CISH^high^ (above median) and CISH^low^ (below median), we found that the CISH^high^ subset exhibited significantly lower expression of *41BB* and *PDCD1*, while CISH^low^ cells had high levels of *41BB* and *PDCD1*.

In order to determine the functional relevance of increased PD1 expression in the absence of CISH we employed the pmel-1 *Cish*^-/-^ murine melanoma model that uses a TCR transgenic which recognizes both murine and human melanoma antigen gp100^15^. T cells from WT and *Cish* KO pmel-1 congenically marked thy1.1 splenocytes were enriched and adoptively transferred into syngeneic C57BL/6 bearing B16 melanomas that express their antigen hgp100. Eight days post ACT, T cells were extracted from tumors, and evaluated for PD1 and the congenic marker thy1.1 expression by flow cytometry. Here, we found that PD1 was expressed significantly higher in *Cish* KO T cells compared to wildtype littermates (**Fig. 6c**). This observation suggested that *Cish* KO T cell therapy would further benefit from PD1 blockade. Using the same ACT approach, a subset of mice also received anti-PD1 or isotype control as indicated. Untreated mice, mice receiving isotype antibody or mice receiving anti-PD1 alone quickly succumbed to their tumors (**Fig. 6d**). The ACT of WT *Cish* pmel-1 T cells conferred only a marginal improvement in tumor treatment, whereas both *Cish* KO or combination of pmel-1 WT with anti-PD1 antibody significantly slowed tumor growth compared to anti-PD1 alone. The combination of pmel-1 *Cish* KO and anti-PD1 resulted in profound tumor regression. Moreover, the combination of *Cish* KO T cells and anti-PD1 treatment resulted in long-term survival (**Fig. 6e**). These findings indicate that inhibition of CISH may improve the outcome of PD1 inhibition and that unleashing the internal and external potential of neoantigen T cells may greatly enhance the effectiveness of adoptive immunotherapy.

## Discussion

Cancer immunotherapy offers the potential of using a patient’s own immune cells to mediate the destruction of metastatic cancer. Residing in the patient’s own lesions are tumor-specific T cells that have the specificity for cancer cells. Despite the presence of these specific T cells, in many cases cancer growth continues unabated^46^. Adoptive transfer of these *ex vivo* stimulated TIL can result in tumor regression and extended survival in patients with melanoma^12^. Yet despite the promise of TIL therapy, and despite the abundance of TIL that are accessible within the tumor microenvironment, a significant proportion of TIL fail to elicit durable tumor regression. The mechanisms by which tumor microenvironments (TME) suppress T cell function are multifaceted and complex. Low antigen density, and reduced MHC expression by tumor cells and more recently cell surface immune checkpoint engagement, are among the core extrinsic contributors to unproductive responses to TCR ligation, preventing the initiation of potent and sustained cytolytic responses by tumor resident T cells. Checkpoint inhibition has improved regression of metastatic melanoma^47-49^, underscoring the value of targeted approaches to increase and prolong the antigen-specific T cell response. Comparatively, the identification and therapeutic targeting of important T cell intrinsic checkpoint mechanisms is far less developed, owing in part to the previous inherent difficulties in drugging intracellular target proteins.

T cells within the tumor microenvironment have been reported to be in a constant state of chronic state of antigenic stimulation and express markers of activation/exhaustion such as CD39, 41BB, TOX and PD1^19,50^. Surprisingly, we observed that CISH was inversely expressed with these established activation/exhaustion markers in tumor resident T cells. CISH expression was enriched in tumor-resident T cells and found to be induced by TCR-specific stimulation with maximal expression positively correlating with T cell maturation status. We sought to explore the physiological importance of *CISH* in human T cells present in the TME.

While the inactivation and blockade of classical cell-surface immune checkpoint targets such as PD-1 is readily achievable through monoclonal antibody-based targeting, intracellular signaling molecules such as CISH have historically been difficult to modulate with comparable efficacy and specificity. The advent of targeted gene editing tools such as CRISPR/Cas9 have allowed for precise, efficient, and permanent gene inactivation in human T cells^25^. In the current study we developed a CRISPR/Cas9 editing strategy that enables efficient and precise genetic disruption of *CISH* in human T cells and used this capability to characterize the impact of *CISH* KO on T cell function. Furthermore, we have been able to successfully adapt our CRISPR/Cas9 approach to a clinical scale, cGMP-compatible production of *CISH* KO neoantigen specific TIL to achieve highly efficient (>95%) and reproducible *CISH* KO in T cells derived from tumor fragments from multiple patients. Remarkably, NGS and GUIDE-seq analysis of CRISPR editing specificity identified no measurable OT activity in *CISH* KO T cells, leading us to conclude the CRISPR editing of CISH in patient TIL was specific with minimal risk of genotoxic events.

*CISH* KO human T cells exhibited enhanced cytokine production; both in magnitude and polyfunctionality after TCR cross-linking. In addition, transduction of TCRs specific to cancer-testis antigen NY-ESO-1 coupled with CISH deletion resulted in profound increase in specific effector cytokine production and relevant tumor killing. Functional assessment after CISH deletion in TIL from three GI cancer patients revealed a consistent enhancement in mutant neoantigen reactivity and that was not observed against naturally found wildtype antigens. This was effector cytokine response was maximally higher (in some cases 5-fold more) and retained over several antigen dose titrations. This indicates that *CISH* inhibition not only increases the overall functionality of TIL but also enhances sensitivity to antigenic stimulation. The increase in antigen avidity could be of critical value in TIL immunotherapy where tumors likely express low levels of neoantigen. Furthermore, neoantigen reactivity is often ‘lost’ after rapid expansion prior to ACT. We found that inactivation of CISH in patient TIL restored the neoantigen reactivity that was lost in control TIL after traditional TIL expansion resulting in a significant increase in TIL functionality against a tumor neoantigen. Overall, these data imply that *CISH* KO increases TCR functional avidity of TIL to mutation-specific neoantigens.

Advances in DNA sequencing has made personalized precision medicine an attainable possibility^7^. At present, however, there remain many logistical challenges facing neoantigen TIL therapy. Access to universal hotspot mutations could obviate many of these limitations if reactivities could be consistently found in patients^51^. We found an enrichment of *CISH* expression in tumor resections lacking reactivity to hotspot *TP53* mutations. CISH *KO* did not alter tetramer-positive precursor frequency but did significantly enhance reactivity against mutant p53. The deletion of CISH in conjunction with the transduction of a TCR library targeting shared hotspot mutations and HLAs could enable the broad application of neoantigen adoptive immunotherapy.

In our study we found that the *CISH* KO in mature patient TIL not only unleashes neoantigen reactivity, but also initiates a hyper-activation program with a significant increase in metabolic activity and expansion. We and others have observed that increased activation is often associated with progressive maturation and functional *in vivo* senecence^52,53^ and observed that AKT signaling plays a key role in that process^44,54^. Molecular dissection of these activation pathways revealed that CISH selectively inhibited PLCγ1. Subsequently, the deletion of CISH resulted in increased ERK, NFAT and NFAT activity but not AKT in human T cells. This selective uncoupling of activation and maturation by CISH deletion might give insights into these critical biological processes and potential therapies.

TIL express a number of characterized activation/exhaustion markers that have been the focus of much study^13,55^. We were surprised to observe that CISH was largely inversely expressed with these markers in minimally processed tumor derivied T cells. Analysis of scRNAseq of CISH depleted T cells revealed that CISH-low cells had high levels of PD1 and in preclinical model the tumor-specific Cish-deficient T cells had in an increase in PD1 expression in the tumor. This increased PD1 expression in the absence of CISH raised the question whether the inhibition of PD-1 in CISH-deficient T cells could further enhance their anti-tumor responses. We addressed this question using a pmel-1 Cish^-/-^ murine melanoma model that uses a TCR transgenic which recognizes both murine and human melanoma antigen gp100. We found that the transfer of CISH-deficient T cells in combination with anti-PD-1 antibody resulted in a significant and synergistic response, leading to a significant and durable tumor regression and complete survival of tumor-bearing mice out to more than 70 days. These findings indicate that removing or blocking the intrinsic suppression of the of neoantigen reactivity of T cells via both CISH and PD-1 checkpoint blockade may significantly enhance the efficacy of adoptive cell immunotherapy in the clinical setting.

Our findings demonstrate that advances in gene editing can result in successful modification of fully mature TIL and reverse internal suppression of the of neoantigen reactivity, making them more functional, proliferative and susceptible to checkpoint blockade. We demonstrate that CISH inhibition can improve the anti-tumor responses of T cells and the ability to efficiently and precisely inactivate CISH using CRISPR, positions CISH as a next-generation immune checkpoint target that may enhance clinical efficacy in the setting of solid tumors. To this end, these pre-clinical data are the foundation for a recently initiated human clinical trial entitled “A Study of Metastatic Gastrointestinal Cancers Treated with Tumor Infiltrating Lymphocytes in Which the Gene Encoding the Intracellular Immune Checkpoint CISH Is Inhibited Using CRISPR Genetic Engineering” (ClinicalTrials.gov Identifier NCT04426669). Results from this trial will hopefully shed light on the feasibility, safety and efficacy of novel checkpoint inhibition using neoantigen-selected, CRISPR genetically engineered *CISH* KO T cell therapy for solid tumors.

## Methods

### Study approval

Animal experiments were conducted with the approval of the NCI Animal Use and Care Committees and performed in accordance with NIH guidelines. All NIH volunteers and patients providing human samples were enrolled in clinical trials approved by the NIH Clinical Center and NCI institutional review boards. Each patient signed an informed consent form and received a patient information form before participation.

### Mice and cell lines

C57BL/6 mice (obtained from Charles River Laboratories, Frederick, MD) of 6–8 weeks of age were used as recipient hosts for adoptive transfer unless otherwise indicated. pmel thy1.1 transgenic mice (B6.Cg-/Cy Tg [TcraTcrb] 8Rest/J) were used for adoptive cell transfer experiments. All mice were maintained under specific pathogen-free conditions. Modified B16-mhgp100 (H-2Db), a mouse melanoma cell line, was transduced as previously described to express glycoprotein 100 (gp100) mutated to express human amino acid residues at positions 25–27 (EGS to KVP); this line was used as the tumor model. Cell lines were maintained in complete media DMEM (Gibco) with 10% FBS, 2-Mercaptoethanol, 1% glutamine and 1% penicillin–streptomycin.

### Immunoblot analysis

Western blot analysis was performed using standard protocols. Proteins were separated by 4%–12% SDS-PAGE, followed by standard immunoblot analysis using anti–CISH and β-actin (Cell Signaling). In brief, for immunoblot quantifications, cells were resuspended in total cell extraction buffer and kept on ice for 10 min followed by homogenization. Cells were then centrifuged at 20,000g for 20 min at 4°C to pellet cell debris. Detection of proteins was performed using secondary antibodies conjugated to horseradish peroxidase-HRP and the super signal west pico chemiluminescent substrate (Thermo Scientific-Pierce).

### Retroviral transduction

To produce the γ-retrovirus, package cell line 293GP were cotransfected with 9□µg of target vector DNA and 4□µg envelope plasmid (RD114 envelope was used to produce virus to infect human T cells; pEco envelope was used to produce virus to infect murine T cells) using lipofectamine 2000 (Cat. No. 11668019, Invitrogen, Carlsbad, California, USA) on a 100□mm^2^ poly-D-lysine–coated plate (Corning, New York, USA). Viral supernatants were harvested 48 and 72□hours after transfection. For T-cell transduction, human peripheral blood mononuclear cells were activated with 50□ng/mL OKT3 (Cat. No. 130-093-387, Miltenyi Biotec) and harvested for retroviral transduction on day 2. Cells were applied to vector-preloaded RetroNectin (Takara) coated non-tissue culture 6-well plates (Corning) at a concentration of 1×10^6^ per well and centrifuged at 1500rpm at 32°C for 10 minutes. After centrifugation, the cells were then cultured in AIM-V medium containing 10% human AB serum (Valley Biomedical) and 300□IU/mL IL-2 until use.

### Peripheral blood T cell editing with CRISPR/Cas9

PBL T cells were stimulated using anti-CD3/CD28 dynabeads in X-Vivo 15 supplemented with 10% human AB serum, 300IU/ml IL-2, and 5 ng/mL IL-7 and IL-15 for 48 hours prior to electroporation. T cells were electroporated with 15ug Cas9 mRNA and 10ug CISH sgRNA using the Neon electroporator (3e6 in 100ul tip) and pulse conditions 1400V, 10ms, 3 pulses. Electroporated T cells were recovered in T cell media without antibiotics for 30min before bringing to 1e6/ml in complete T cell media.

### Production of *CISH* KO TIL using CRISPR/Cas9

Interleukin-2 expanded tumor infiltrating lymphocytes (TIL) were thawed (day −5) and allowed to recover for 24 hours in TIL medium (X-Vivo 15, 10% human AB serum, 6000 IU/mL IL-2, and 5 ng/mL IL-7 and IL-15) at 37°C, 5% CO2. After the initial rest period (day −4), TIL cultures were harvested and a volume reduction step was performed prior to re-suspenion in fresh TIL media followed by stimulation with plate bound (5 µg/ml) anti-CD3 (OKT3) and soluble anti-CD28 (2 µg/mL) for 4 days at 37°C, 5% CO2. Four days later (day 0), stimulated TIL were washed with PBS and re-suspended at 2.5 x 10^7^ NC/mL in either PBS (GMP process) or Neon buffer T. Each 2.5 x 10^6^ viable TIL were electroporated with 15µg Cas9 mRNA and 10µg *CISH* gRNA in a 100µl tip using the Neon electroporation device (Life Technologies) using parameters 1400v, 10 ms width, 3 pulses. For non-REP expansion, TIL were immediately returned to TIL medium and maintained at ∼1 x 10^6^ viable cells/ml with either media addition or 50% volume exchange as required.

### Rapid expansion of *CISH KO* TIL

For rapid expansion protocol (REP), electroporated TIL were immediately transferred to TIL REP media (X-Vivo 15, 5% human AB serum, 3000 IU/mL IL-2) and seeded at 5 – 7.5 x 10^3^ viable TIL per cm^2^ in G-Rex culture vessels (Wilson Wolf, New Brighton MN) and combined with either autologous or allogeneic (3 pooled donors) irradiated PBMC feeders at a ratio of 1 TIL to 100 feeders (1:100). G-Rex vessels were incubated for 6-8 days at 37°C, 5% CO2. On day 6-8, the culture was evaluated and split according to the following: if viable NC/mL < 1 x 10^6^ VNC/mL, a 1:3 split was performed; if viable NC/mL > 1 x 10^6^ VNC/mL, a 1:6 split was performed. Each G-Rex was equally transferred to 2 or 5 additional vessels according to split criteria above and fresh expansion media was added. All vessels were incubated for an additional 6-8 days at 37°C, 5% CO2.

### Neoantigen screening

Tumors from cancer patients were surgically resected at the NIH Clinical Center and subjected to whole-exomic sequencing to identify non-synonymous somatic mutations. TIL cultures derived from individual tumor fragments from a single metastatic colon lesion were initially screened for reactivity against multiple TMG constructs or peptide pools using the enzyme-linked immunospot (ELISPOT) assay and flow cytometric evaluation of up-regulation of the T cell activation marker 4-1BB as previously described^7^.

### Intracellular Cytokine Staining (ICS)

To evaluate the functionality of *CISH* KO T cells they were assayed for specific release of functional cytokines in a co-culture with neoantigen loaded APCs. APCs, either B cells or DCs as indicated, were generated from autologous PBMCs and cultured for 5 days. Concurrently, the cryo-preserved gene-modified cells from were thawed in pre-warmed complete media supplemented with IL-2 (300 IU/mL) and grown for 2 days. On the day of the co-culture, APCs were pulsed with mutant or wildtype (WT) peptides for 2 hours and then washed prior to being mixed with either *CISH* KO T cells or Control T cells. The co-culture was setup in the presence of golgi-blocking reagents and allowed to continue for 6 hours. After the 6 hours, samples were extracellular stained for T cell makers, fixed and permeabilized, then stained for the cytokines IFN-γ, IL-2 and TNF-α. Flow cytometry was then performed, and cells were analyzed for specific function.

### Cytotoxicity

Cytotoxicity assays were carried out with the IncuCyte S3-Platform (Essen BioScience). Adherent 526 (HLA-A2^+^NY-ESO-1^-^) or 624 (HLA-A2^+^NY-ESO-1^+^) tumor cells were plated at 1 × 10^4^ cells per well and incubated overnight at 37°C/5% CO2 in RPMI-1640 medium supplemented with 10% heat-inactivated FBS and GlutaMAX (Life Technologies) in a 96-well flat-bottom plate. The next day, cells were washed and incubated with indicated numbers of NY-ESO-1 TCR-transduced T cells from either control or CISH KO and 3.3 μmol/L IncuCyte Caspase-3/7 reagent (Essen BioScience). Cells were imaged at times indicated to detect apoptosis. Data were analyzed using IncuCyte S3 software (Essen BioScience) to distinguish apoptotic tumor cells from apoptotic T cells.

### GUIDE-seq

GUIDE-seq analysis was performed as described previously using a 6-mismatch cutoff (1), with the following modifications for application to primary lymphocytes. PBL T cells and TIL were stimulated using anti-CD3/CD28 dynabeads for 36-48 hours prior to electroporation in T cell media supplemented with IL-2 (300 IU/ml for PBL-T, and 3000IU/ml for TIL), 5ng/ml IL-7, and 5ng/ml IL-15. Cells were electroporated with 15ug Cas9 mRNA, 10ug CISH sgRNA, and 8-16pmol of GUIDE-seq dsODN using the Neon electroporator at 3e6/100ul tip and pulse conditions 1400V, 10ms, 3 pulses. On-target integration of dsDNA oligo was confirmed by PCR and TIDE analysis prior to NGS.

### GSEA

Gene set enrichment was analyzed using GSEA software (http://software.broadinstitute.org/gsea/downloads.jsp)^56^. Pathway Analysis was performed on the identified differentially expressed genes list using the Core Analysis function included in Ingenuity Pathway Analysis (IPA, Qiagen).

### scRNAseq Capture and library preparation

Single cell suspensions were prepared for single cell RNA-Seq partitioning, barcoding and library generation on the 10x Genomics Chromium platform. Suspensions were washed twice by pelleting cells with centrifugation at 300g in a chilled spinning bucket centrifuge and gentle resuspension in fresh ice-cold PBS with 0.04% BSA. Cell concentrations and viability were determined on a LunaFL fluorescent cell counter using Acridine Orange and Propidium Iodide dye (Logo Biosystems). Suspension concentrations were adjusted and loaded onto the Chromium microfluidic chip using the 3’ v3 gene expression chemistry to target 6,000 barcoded cells, and in samples where fewer cells were available, at the full concentration. Reverse transcription, cDNA amplification, and sequencing library preparation was all performed according to vendor’s user guide.

### Sequencing and primary data processing

Sequencing of final single cell RNA-Seq libraries was performed with the NCI-CCR Genomics Core on the NextSeq 500 platform using 150-cycle v2.5 High Output reagents with a 26bp read for identifying cell-specific barcode and UMI sequences, a 8bp index read for multiplexed sample identity, and a 98bp read to identify the cDNA insert. Multiplexed samples were sequenced multiple times to achieve target read depth. Data was processed with the 10x Genomics cellranger v3.0.1 pipeline to generate sample fastq sets followed by alignment of reads to the human GRCh38 reference sequence prepared by 10x Genomics (refdata-cellranger-GRCh38-3.0.0), generation of a cell barcode by expressed gene matrix, and basic quality metrics of capture, library and sequencing performance.

### Supported lipid bilayer (SLB) and calcium flux measurement

Liposomes containing 6.25% DGS-NTA and 2% Cap-Biotin-DOPC lipids were prepared using an extruder method as per manufacturer’s instruction (Avanti Polar Lipids, Inc.). Briefly, liposomes were applied to charged cover glasses for about 30 to form planar bilayer. The lipid bilayer was washed with HBS BSA buffer and then incubated with streptavidin (5mg/ml) for 30 min at RT and then with mono-biotinylated anti-CD3 antibody (1µg/ml OKT3, eBiosciences), histidine tagged CD80 (100 molecules/µm^2^), and ICAM-1 (100 molecules/µm^2^). Finally, the SLB was washed and brought to 37°C before injecting the cells. Calcium flux measurement was done according to the protocol as described by Skokos et al 2007. WT or CISH KO T cells were labeled with 2μM Fura-2-Am in serum free HBS buffer for 30 min at room temperature followed by de-esterification of the dye for another 30 min at 37°C in serum containing buffer. Imaging of cells was performed using 40X 1.35NA UApo 340 objective. Images of cells were acquired at a distance of 3 mm from the interference reflection microscopy image plane to acquire fluorescence from the equatorial plane of the cell. Cells were imaged live at 37°C while interacting with the SLB in HBS buffer containing Ca^2+^ and Mg^2+^. The Fura-2 emission at 510 nm upon excitation with both 340 and 380 nm light was captured for a few fields of cells. These fields of cells were repeatedly imaged for 40 min to obtain a time course of multiple cells. At the end of the experiment cells were treated with a buffer containing 1 μM Ionomycin, 20 mM Calcium and 2mM Magnesium to record the high calcium condition followed by a treatment with a buffer containing 3mM Magnesium, 2mM EGTA and no calcium to record the low calcium condition. Image analysis was performed using Metamorph software.

### Metabolomics

T cells from 6 donors were flash frozen before or after 4 hours of stimulation with CD3 crosslinking. Samples were then analyzed for metabolites using Metabolons inhouse services. The present human dataset comprises a total of 251 compounds of known identity (named biochemicals). Following normalization to DNA concentration, log transformation and imputation of missing values, if any, with the minimum observed value for each compound, Paired *t*-tests and Welch’s two-sample *t*-test were used to identify biochemicals that differed significantly between experimental groups. For OCR, T cells were measured at 37°C using an Xfe96 extracellular analyzer (Seahorse Bioscience). Briefly, 10^6^ cultured T cells were initially plated on poly-L-Lysine coated XF96 well plate in unbuffered DMEM (DMEM with 25 mM glucose as indicated, 1 mM sodium pyruvate,32 mM NaCl, 2 mM GlutaMax, pH 7.4) and incubated in a non-CO2 incubator for 30 minutes at 37°C. OCR was calculated using Seahorse XFe-96 proprietary software.

### Statistics

Significance was determined by either student *t* test or ANOVA for repeated measures, *P >0.05, **P>0.01, ***P>0.001, ****P>0.0001.

## Extended Data

**Extended Data Fig. 1. Optimization of CISH KO by CRISPR**.

**a**. Western blot analysis after CISH editing by several different individual modified guide-RNAs and Cas9. **b**. Intracellular cytokine staining after CISH deletion and TCR crosslinking.

**Extended Data Fig. 2. Molecular characterization of CISH KO in TIL**

**a**. Schematic of modified cGMP rapid expansion protocol (REP) used for CRISPR/Cas9 editing of *CISH* in human tumor infiltrating lymphocytes. **b**. TiDE analysis of InDels after CISH editing in TIL. **c**. Western blot analysis after CISH deletion in multiple donors. **d**. Total fold-expansion of control and CISH KO TIL during 14 day REP manufacturing process. **e**. GUIDE-Seq output following NGS and sequence read analysis. On target read number is indicated to the right of the sequence alignment. **f**. TIL expansion in long-term (21 day) cultures after CISH editing and REP with or without IL-2.

**Extended Table 1. Computationally predicted off-target loci evaluated by targeted-amplicon sequencing**.

**Extended Data Fig. 3. Screening for potential neoantigens in patient A**.

The tumor is split after excision, in one fragment T cells are grown out and in the other fragment the tumor is subjected to exomic and RNA sequencing. Non-synonymous point-mutations predicted to bind to autologous MHC are made into tandem mini genes or peptides targets. T cells derived from the tumor resection and neoantigen loaded autologous APCs are then co-cultured and assayed for the upregulation of IFN-γ or 41BB.

**Extended Data Fig. 4. Screening for potential neoantigens in patient B**.

**a**. Confirmation of specific neoantigen recognition. **b**. Flow cytometric analysis of neoantigen specific TIL.

**Extended Data Fig. 5. RNA-SEQ resections with P53 hotspots**

**a**. PCA analysis of RNA-SEQ resections with P53 hotspots that were found to have or not have p53 reactivities. **b**. GSEA profiling of RNA-SEQ resections with P53 hotspots that were found to have p53 reactivities.

**Extended Data Fig. 6. Metabolic analysis of CISH KO T cells**.

a, 10 days after stimulation of and deletion of CISH CD8+ enriched T cells from PBL were subjected to metabolite analysis with or without a 4-hour CD3 stimulation. All samples were normalized with benchmarked metabolites and normalized to DNA content. Standard statistical analyses are performed at Metabolon using ArrayStudio on log transformed data and Welch’s two-sample t-test. **b**, OCR of CISH replete or CISH KO TIL in response to indicated metabolic modulators. Representative of two independent patients. Statistical significance was determined by either student *t* test or ANOVA for repeated measures, *P >0.05, **P>0.01, ***P>0.001, ****P>0.0001. Error bars denotes +/-SD.

## Acknowledgements

Support provided, in part through the Intramural program CCR at the National Cancer Institute. Support from CCR Single Cell Analysis Facility was funded by FNLCR Contract HSN261200800001E. Sequencing was performed with the CCR Genomics Core. This work utilized the computational resources of the NIH HPC Biowulf cluster. (http://hpc.nih.gov) Next generation sequencing and bioinformatic analysis of CRISPR off-target activity was conducted at the University of Minnesota Genomics Center (UMGC).

## Contributions

D.C.P., B.R.W., S.A.R., B.S.M. and N.P.R. were involved in study design. M.C., S.A.R., B.S.M., P.F.R., R.S.M. contributed to study concepts. B.R.W., D.S., D.H.M., N.J.S., M.D.D. were responsible for manufacturing and validating therapeutic cells. D.C.P., B.R.W., Y.P., C.M.K., F.J.L., R.J.K., D.G., Z.F., S.K.V., K.P., P.M., D.C.D., S.J.P., D.S., N.J.S., M.D.D., M.J.J., W.S.L., N.K.W., L.V., T.H., T.B., O.B. were involved in data acquisition. J.J.G., T.D.P. were involved in quality control of data and algorithms. Y.H., C.Y., D. M., L.J. were involved in data analysis and interpretation.D.C.P., B.R.W, Y.H., L.J. contributed to statistical analysis. D.C.P. and B.R.W. wrote the manuscript. All authors approved the article for submission and publication.

## Conflicts of Interest

M.C. is a co-founder of Intima Bioscience. B.R.W, S.A.R, and B.S.M have received sponsored research support from Intima Bioscience. D.C.P., B.R. W, M.C., S.A.R., B.S.M, and N.P.R. have patents filed based on the findings described here. The authors declare no competing interests.

