## Extended data for "Internal checkpoint regulates T cell neoantigen reactivity and susceptibility to PD1 blockade"

### Slide 1
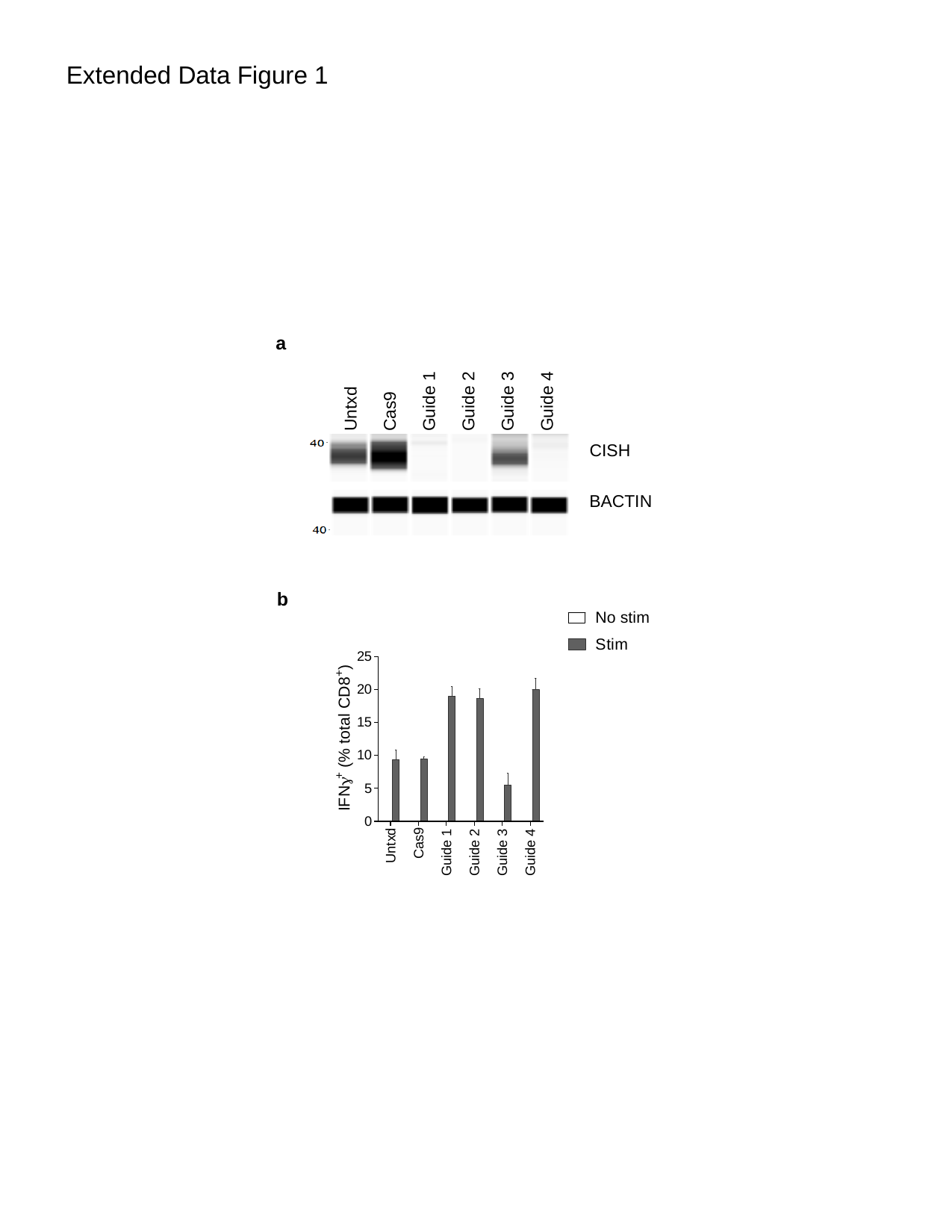

Extended Data Figure 1
a
Guide 4
Untxd
Cas9
Guide 1
Guide 2
Guide 3
CISH
BACTIN
b

### Slide 2
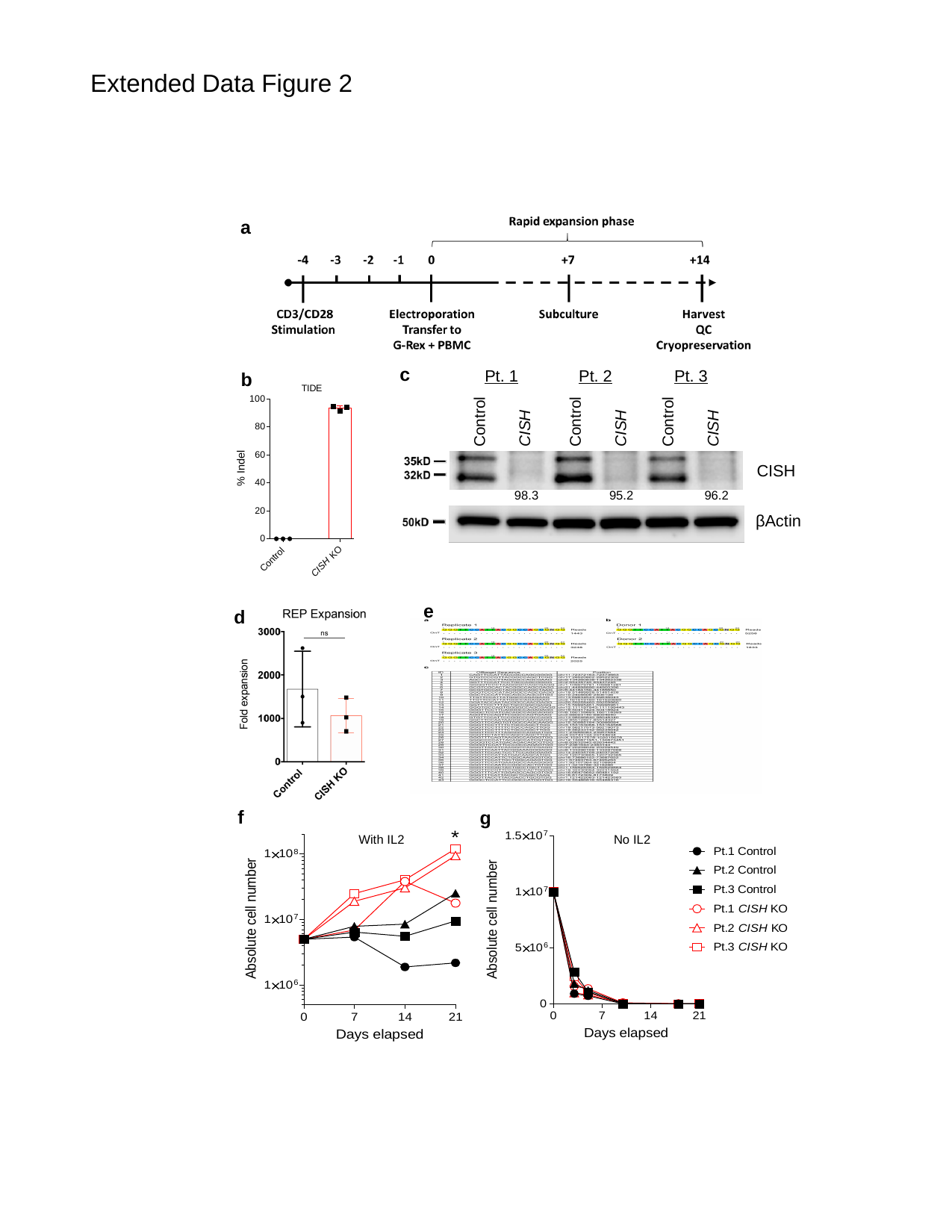

Extended Data Figure 2
a
c
Pt. 1
Pt. 2
Pt. 3
Control
Control
Control
CISH
CISH
CISH
CISH
98.3
95.2
96.2
βActin
b
e
d
f
g
With IL2
No IL2

### Slide 3
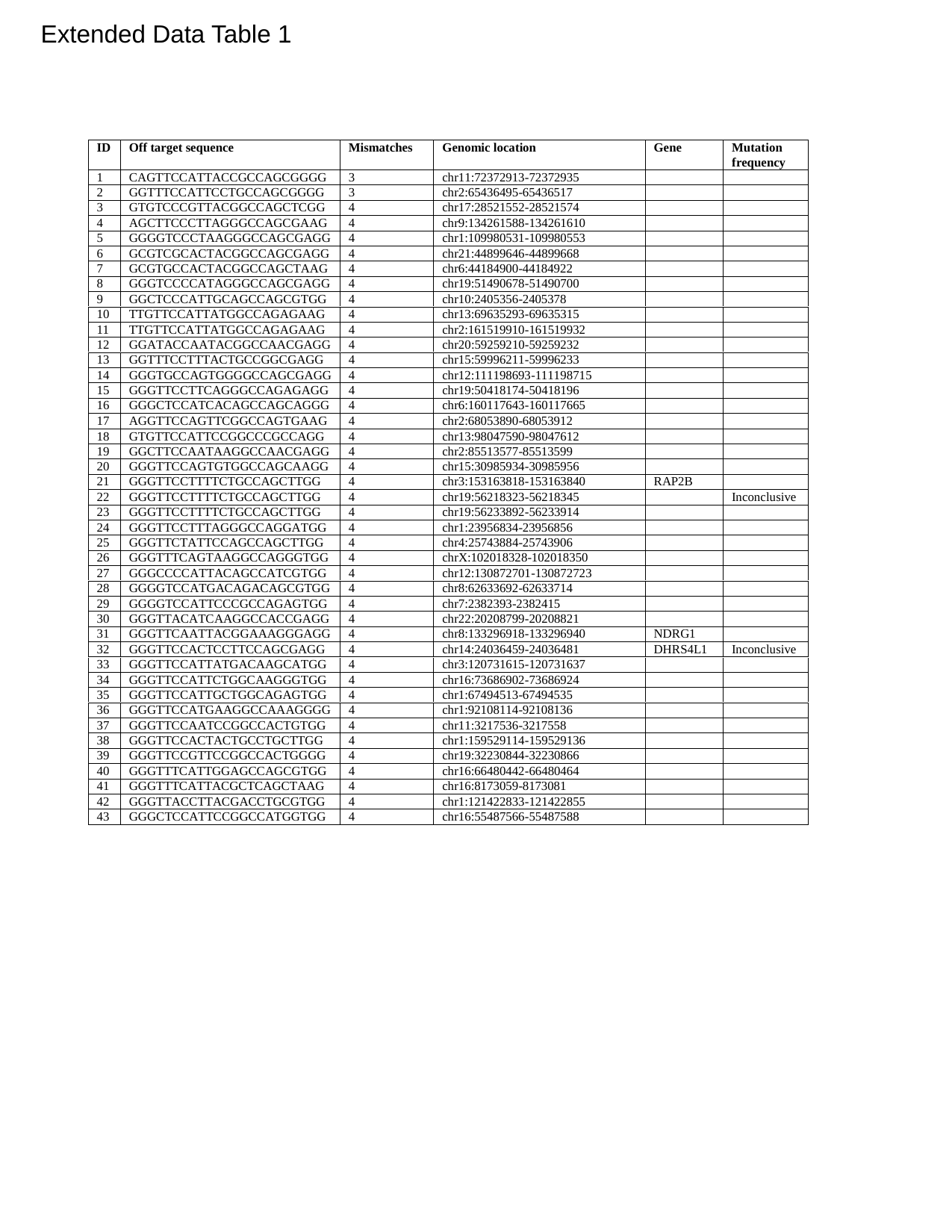

Extended Data Table 1

### Slide 4
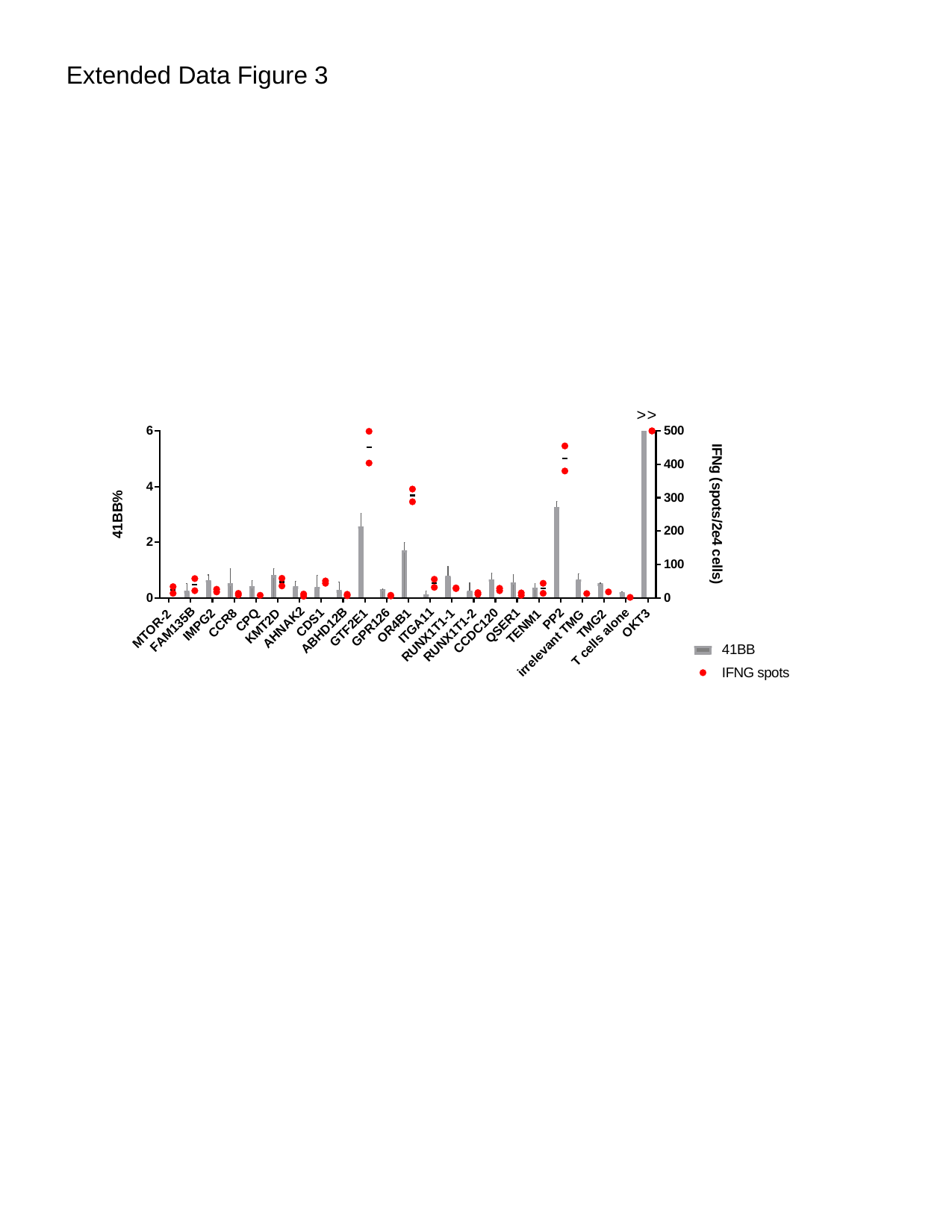

Extended Data Figure 3
>
>

### Slide 5
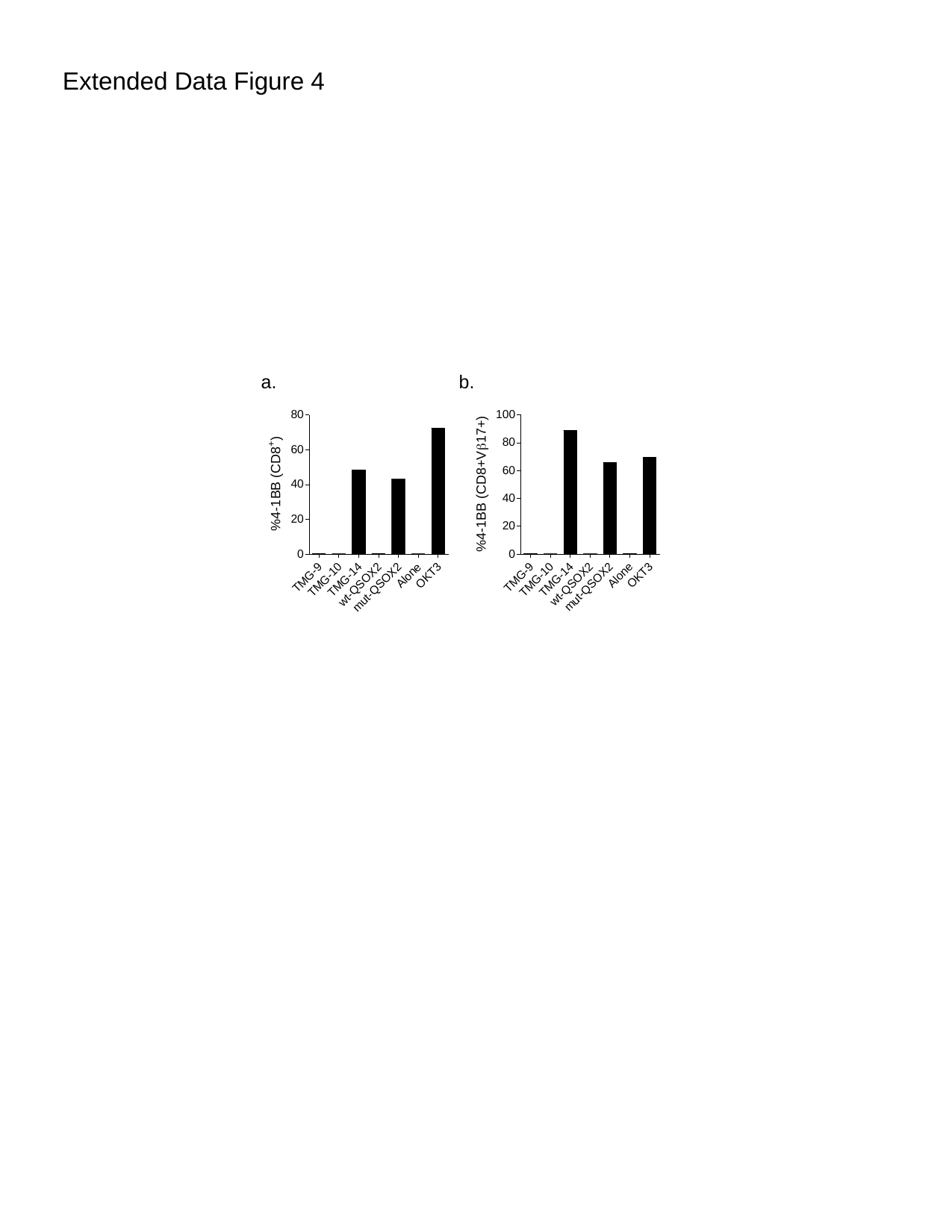

Extended Data Figure 4
a.
b.

### Slide 6
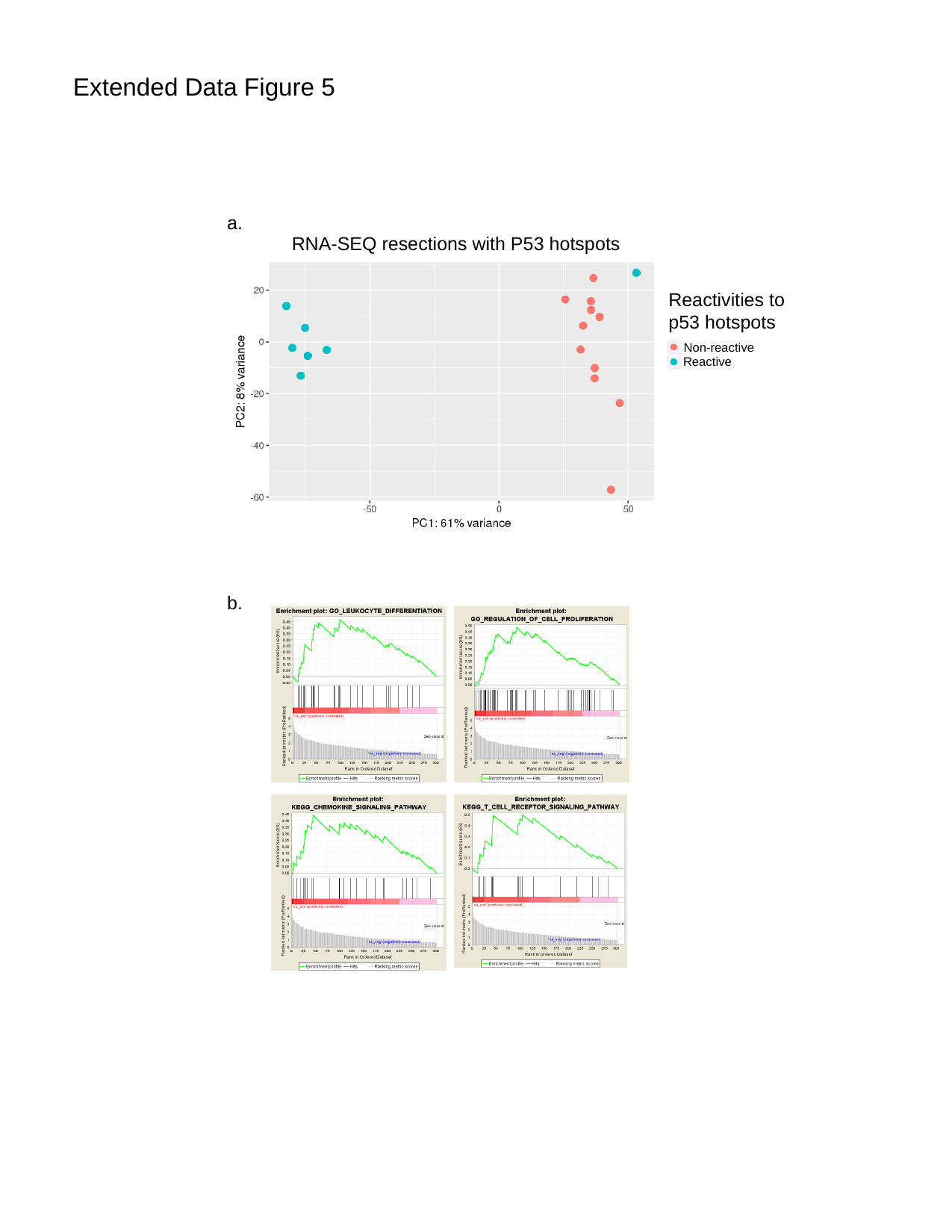

Extended Data Figure 5
a.
RNA-SEQ resections with P53 hotspots
Reactivities to p53 hotspots
Non-reactive
Reactive
b.

### Slide 7
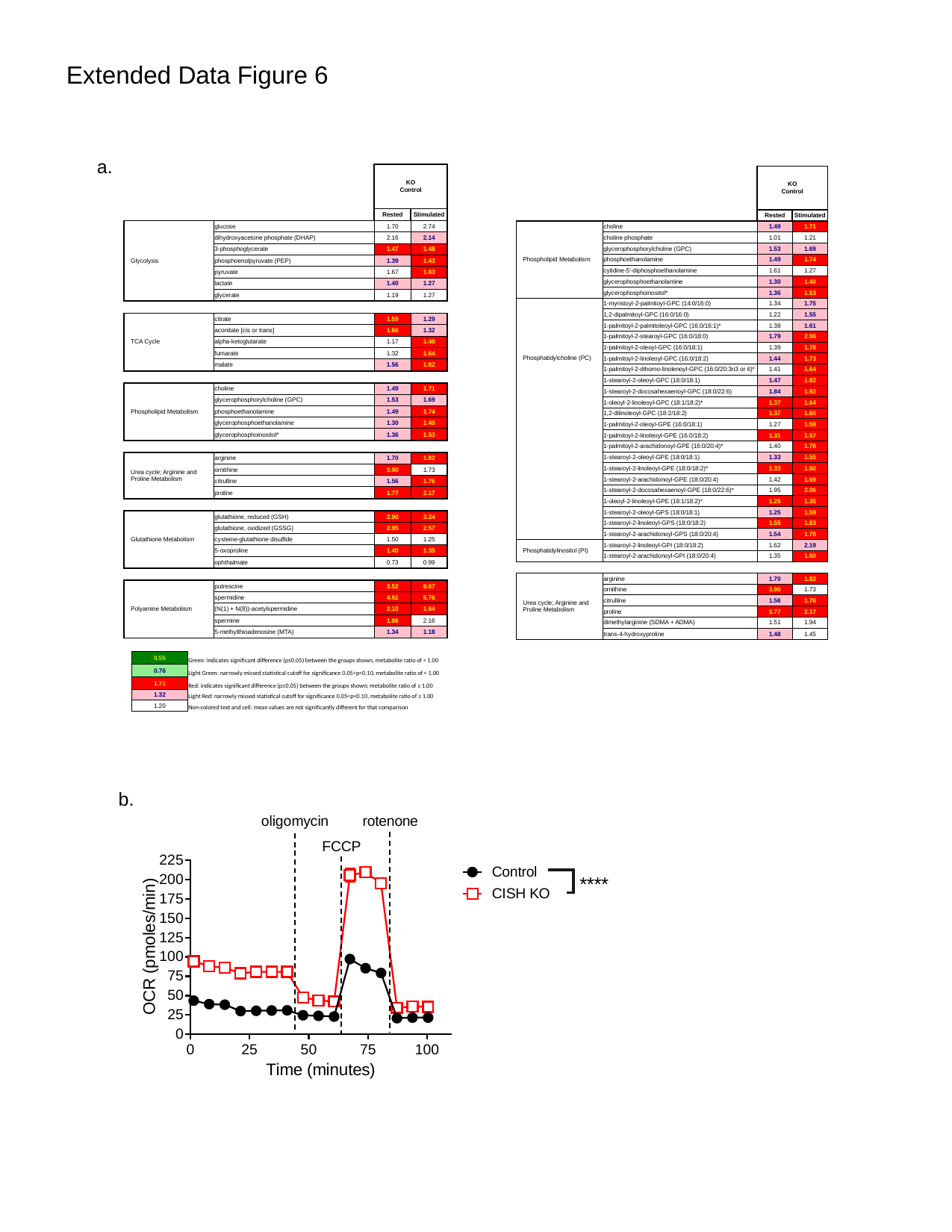

Extended Data Figure 6
a.
| | | KO Control | |
| --- | --- | --- | --- |
| | | Rested | Stimulated |
| Glycolysis | glucose | 1.70 | 2.74 |
| | dihydroxyacetone phosphate (DHAP) | 2.16 | 2.14 |
| | 3-phosphoglycerate | 1.47 | 1.48 |
| | phosphoenolpyruvate (PEP) | 1.39 | 1.43 |
| | pyruvate | 1.67 | 1.83 |
| | lactate | 1.40 | 1.27 |
| | glycerate | 1.19 | 1.27 |
| TCA Cycle | citrate | 1.59 | 1.29 |
| | aconitate [cis or trans] | 1.66 | 1.32 |
| | alpha-ketoglutarate | 1.17 | 1.40 |
| | fumarate | 1.32 | 1.64 |
| | malate | 1.56 | 1.82 |
| Phospholipid Metabolism | choline | 1.49 | 1.71 |
| | glycerophosphorylcholine (GPC) | 1.53 | 1.69 |
| | phosphoethanolamine | 1.49 | 1.74 |
| | glycerophosphoethanolamine | 1.30 | 1.40 |
| | glycerophosphoinositol\* | 1.36 | 1.53 |
| Urea cycle; Arginine and Proline Metabolism | arginine | 1.70 | 1.82 |
| | ornithine | 3.90 | 1.73 |
| | citrulline | 1.56 | 1.76 |
| | proline | 1.77 | 2.17 |
| Glutathione Metabolism | glutathione, reduced (GSH) | 2.90 | 3.24 |
| | glutathione, oxidized (GSSG) | 2.95 | 2.57 |
| | cysteine-glutathione disulfide | 1.50 | 1.25 |
| | 5-oxoproline | 1.40 | 1.35 |
| | ophthalmate | 0.73 | 0.99 |
| Polyamine Metabolism | putrescine | 3.52 | 8.67 |
| | spermidine | 4.92 | 5.76 |
| | (N(1) + N(8))-acetylspermidine | 2.10 | 1.64 |
| | spermine | 1.96 | 2.16 |
| | 5-methylthioadenosine (MTA) | 1.34 | 1.18 |
| | | KO Control | |
| --- | --- | --- | --- |
| | | Rested | Stimulated |
| Phospholipid Metabolism | choline | 1.49 | 1.71 |
| | choline phosphate | 1.01 | 1.21 |
| | glycerophosphorylcholine (GPC) | 1.53 | 1.69 |
| | phosphoethanolamine | 1.49 | 1.74 |
| | cytidine-5'-diphosphoethanolamine | 1.61 | 1.27 |
| | glycerophosphoethanolamine | 1.30 | 1.40 |
| | glycerophosphoinositol\* | 1.36 | 1.53 |
| Phosphatidylcholine (PC) | 1-myristoyl-2-palmitoyl-GPC (14:0/16:0) | 1.34 | 1.75 |
| | 1,2-dipalmitoyl-GPC (16:0/16:0) | 1.22 | 1.55 |
| | 1-palmitoyl-2-palmitoleoyl-GPC (16:0/16:1)\* | 1.38 | 1.61 |
| | 1-palmitoyl-2-stearoyl-GPC (16:0/18:0) | 1.79 | 2.06 |
| | 1-palmitoyl-2-oleoyl-GPC (16:0/18:1) | 1.39 | 1.78 |
| | 1-palmitoyl-2-linoleoyl-GPC (16:0/18:2) | 1.44 | 1.73 |
| | 1-palmitoyl-2-dihomo-linolenoyl-GPC (16:0/20:3n3 or 6)\* | 1.41 | 1.64 |
| | 1-stearoyl-2-oleoyl-GPC (18:0/18:1) | 1.47 | 1.82 |
| | 1-stearoyl-2-docosahexaenoyl-GPC (18:0/22:6) | 1.84 | 1.92 |
| | 1-oleoyl-2-linoleoyl-GPC (18:1/18:2)\* | 1.37 | 1.64 |
| | 1,2-dilinoleoyl-GPC (18:2/18:2) | 1.37 | 1.60 |
| | 1-palmitoyl-2-oleoyl-GPE (16:0/18:1) | 1.27 | 1.59 |
| | 1-palmitoyl-2-linoleoyl-GPE (16:0/18:2) | 1.31 | 1.57 |
| | 1-palmitoyl-2-arachidonoyl-GPE (16:0/20:4)\* | 1.40 | 1.78 |
| | 1-stearoyl-2-oleoyl-GPE (18:0/18:1) | 1.33 | 1.55 |
| | 1-stearoyl-2-linoleoyl-GPE (18:0/18:2)\* | 1.33 | 1.60 |
| | 1-stearoyl-2-arachidonoyl-GPE (18:0/20:4) | 1.42 | 1.69 |
| | 1-stearoyl-2-docosahexaenoyl-GPE (18:0/22:6)\* | 1.95 | 2.06 |
| | 1-oleoyl-2-linoleoyl-GPE (18:1/18:2)\* | 1.25 | 1.35 |
| | 1-stearoyl-2-oleoyl-GPS (18:0/18:1) | 1.25 | 1.59 |
| | 1-stearoyl-2-linoleoyl-GPS (18:0/18:2) | 1.55 | 1.83 |
| | 1-stearoyl-2-arachidonoyl-GPS (18:0/20:4) | 1.54 | 1.78 |
| Phosphatidylinositol (PI) | 1-stearoyl-2-linoleoyl-GPI (18:0/18:2) | 1.62 | 2.19 |
| | 1-stearoyl-2-arachidonoyl-GPI (18:0/20:4) | 1.35 | 1.60 |
| Urea cycle; Arginine and Proline Metabolism | arginine | 1.70 | 1.82 |
| | ornithine | 3.90 | 1.73 |
| | citrulline | 1.56 | 1.76 |
| | proline | 1.77 | 2.17 |
| | dimethylarginine (SDMA + ADMA) | 1.51 | 1.94 |
| | trans-4-hydroxyproline | 1.48 | 1.45 |
| 0.55 | Green: indicates significant difference (p≤0.05) between the groups shown, metabolite ratio of < 1.00 |
| --- | --- |
| 0.76 | Light Green: narrowly missed statistical cutoff for significance 0.05<p<0.10, metabolite ratio of < 1.00 |
| 1.71 | Red: indicates significant difference (p≤0.05) between the groups shown; metabolite ratio of ≥ 1.00 |
| 1.32 | Light Red: narrowly missed statistical cutoff for significance 0.05<p<0.10, metabolite ratio of ≥ 1.00 |
| 1.20 | Non-colored text and cell: mean values are not significantly different for that comparison |
b.
